# Live Mouse Tracker: real-time behavioral analysis of groups of mice

**DOI:** 10.1101/345132

**Authors:** Fabrice de Chaumont, Elodie Ey, Nicolas Torquet, Thibault Lagache, Stéphane Dallongeville, Albane Imbert, Thierry Legou, Anne-Marie Le Sourd, Philippe Faure, Thomas Bourgeron, Jean-Christophe Olivo-Marin

## Abstract

Preclinical studies of psychiatric disorders require the use of animal models to investigate the impact of environmental factors or genetic mutations on complex traits such as decision-making and social interactions. Here, we present a real-time method for behavior analysis of mice housed in groups that couples computer vision, machine learning and Triggered-RFID identification to track and monitor animals over several days in enriched environments. The system extracts a thorough list of individual and collective behavioral traits and provides a unique phenotypic profile for each animal. On mouse models, we study the impact of mutations of genes Shank2 and Shank3 involved in autism. Characterization and integration of data from behavioral profiles of mutated female mice reveals distinctive activity levels and involvement in complex social configuration.

## INTRODUCTION

Mice are routinely used as preclinical model to study the mechanisms leading to human diseases. In the field of psychiatry, assessing mouse social behavior under normal or pathological conditions is critical to understand which neural systems are engaged in these diseases. However, such behaviors are complex and still challenging to investigate in mice given the technical limitations of data gathering and analysis^1^. Indeed, while behavioral protocols investigating one or two animals provide information on activity and cognitive functions such as learning, memory and anxiety, gathering appropriate information on social behaviors requires access to the observation of more than two individuals. Very standardized social interaction tests are available^2,3^ but they rely primarily on quantification of the quality and on the number of simple (dyadic) and short (a few minutes) social interactions, and lack ethologically relevant behavioral markers (e.g., maintenance of social interactions over the long term, social interactions involving more than two mice)^4,5^. In addition, it is necessary to increase the robustness of the data collected and to provide additional extracted features to document thoroughly the dynamics and plasticity of the social aspects over unlimited periods of time for each individual. This is particularly true, for example, for the formation and description of group dynamics of animals.

Solutions to track automatically individuals within a group of mice over hours or days have been proposed (see **Supplementary methods - Motivation and review of the existing tracking methods**). While each of them represents a progress in the field, they all have a number of shortcomings that prevent full exploitation of the data and limit their routine use. One of the main current limitations remains the fact that none of them allow, in the long term and without manual corrections, an individual tracking of a mouse within a group with a sufficient level of details.

To overcome these limitations, we developed a comprehensive system, called *Live Mouse Tracker* (LMT). It allows the automatic live tracking, identification and characterization through behavioral labeling of up to four animals in an enriched environment with no time limit. This solution makes use of RFID sensors and of an infrared/depth RGBD camera, under the control of a machine learning-based framework. RFID provides intermittent identity and tracking validation while the RGBD camera provides continuous depth and volume information^6^ to characterize animal shape and posture (see example with the tracking of one mouse in ^7^). On the basis of this robust central element, we designed a system that can address the whole behavioral workflow and cope with most if not all the difficulties of this type of assays: 1. reproducible acquisition hardware (see **Supplementary methods and movies - Hardware blueprint and assembly instruction; Supplementary PDF file - Assembly instructions**), 2. acquisition calibration (**Supplementary methods - Camera setup and calibration**), 3. standardized data acquisition, 4. live tracking (identity recovery and identity control) (**Supplementary methods - Identity recovery with machine learning, Triggered-RFID closed-loop control**), 5. automatic phenotyping (**Supplementary methods - Behavioral event extraction**), 6. on-the-fly analysis for monitoring the tracking quality or the behavior (**Supplementary methods - Automatic tracking control**), 7. storage of long term observations in database (**Supplementary methods - Database description**), 8. visual data inspection with the development of a specific data player (**Supplementary methods - Database player**) and 9. online data sharing with the community (databases, videos, analysis scripts and results) (**Supplementary methods - Collaborative website sharing data**).

We used LMT with four animals to compare the behavioral profile of female mice lacking *Shank2^8,9^* or *Shank3^8^*, two genes that were found mutated in a subgroup of patients with autism and coding for post-synaptic scaffolding proteins from excitatory synapses^10–13^. We report behavioral differences in the individual profiles between the two strains in the activity levels and involvement in complex social configuration. We confirm that the characteristic differences of activity levels between *Shank2* and *Shank3* mutant mice were still present, even after long interaction periods. Notwithstanding these different activity levels, *Shank2^−/−^* and Shank3^−/−^ mice displayed typical circadian rhythms. We finally show that the atypical social behavior of mutant mice of both strains appeared to disturb the formation of subgroups within mixed-genotype groups of four mice.

These results demonstrate the capability of LMT to study differences in the phenotypes expressed by individuals in groups of mice over larger durations of time.

## RESULTS

We have designed an integrated system that tracks and monitors, for either hours or days, the activities of four mice in a rich environment (**Supplementary movie - Method overview**). This system determines the outline mask and orientation of each mouse and builds, from these data, a comprehensive repertoire of individual and social behavioral events. The system can track mice of any coat color in a rich environment. It is robust to the presence of food, water, sawdust, brown crinkle paper, white compressed cotton cylinders, toys and house (either transparent to infrared or opaque) (**Fig. 1a**). The tracking (**Supplementary Fig. S1 - Tracking diagram**) is performed by using an RGBD camera, filming the mice from the top (**Fig. 1b**). The 2D 1/2 data (infra-red intensity and distance from the sensor of each pixel acquired; **Fig. 1c-d, Supplementary methods - Capturing depth map**) are integrated to compute a background depth map (**Fig. 1e**), which is a representation of the environment where the mice have been computationally removed (**Supplementary methods - Computing the background height-map; Supplementary movie - Background height-map demo**). The segmentation (**Fig. 1f-g**) step that extracts objects and boundaries is performed on an image obtained by subtracting the current acquisition from the background height-map (**Supplementary methods - Segmentation and detection**). Segmentations are then filtered by a dedicated machine learning to reject detections that do not match the mice (**Supplementary methods - Building the detection feature vector, Detection filtering with machine learning**) and detections are then processed to separate mice that are in contact (**Supplementary methods - Detection splitter**). Extraction of additional data such as the orientation (**Supplementary methods - Head/tail detection post-processing**) or the detection of ears, eyes and nose are then performed (**Fig. 1f-g**, **Supplementary methods - Head sub-parts detection**) thus enabling the detection of the tilt-orientation of the head. Detections are then processed for tracking (**Supplementary methods - Tracking extender association process**). Identity of tracks is retrieved in real-time by combining machine learning (**Supplementary methods - Identity recovery with machine learning**) with RFID (**Supplementary methods - RFID calibration, Triggered-RFID closed-loop control, RFID hardware**). Detection, tracking, and RFID readings information are stored in a database (**Supplementary methods - Database description**) and can be investigated live during the tracking (**Supplementary methods - Querying database information with R and Python**), or accessed via our online network (**Supplementary methods - UDP live network information stream**). Video and background maps are simultaneously recorded respectively in movies and image series (**Supplementary methods - MP4 video recording**).

Tracking reliability was assessed on four experiments of ten minutes each (i.e. 18,000 frames each experiment) with respectively one to four mice in the same cage. LMT detected the mice at least 99.25% of the time (i.e. detection rate), for any number of mice (from one to four, **Fig. 1j**). Then two independent experts conducted validations of the detection (**Supplementary methods - Manual validation (extra information) and Fig.S3- 5)** to estimate reliability of 1) segmentation (**Fig. 1k**), essential to determine if animals are in contact and to conduct shape analysis; 2) head-tail orientation (**Fig. 1l**), needed for asymmetrical events; 3) identification of individual mice (**Fig. 1m**), essential to understand inter-individual relationships. Experts used the integrated database player (**Supplementary methods - Database player**) to manually validate frame by frame the tracker performances. In a group of four, mice were correctly segmented (i.e., the mask fit the real body shape exactly, no extra inclusion of objects or reflection on the walls were taken into account, no merging of two animals) in more than 95.75% of the frames in which the mouse was detected (**Fig. 1k**). Detected orientation was accurate in more than 99.36% of the detected frames (**Fig. 1l**). Finally, the identity error rate did not exceed 2.69% for a group of four mice (**Fig. 1m**). Overall, the system keeps track of the identities of the animals, corrects false identifications and prevents any error from propagating thanks to the RFID used in the system. Switching identities episodes have a mean duration of 1.64s ± 0.23 s for the expert #1, and 1.33s ± 0.23 s for the expert #2 (**Supplementary methods - Manual validation (extra information); Supplementary Fig. S3a**). All these estimates allowed us to calculate the Multiple Object Tracking Accuracy (MOTA) tracking performance index^14^, which considers false positive, false negative and identity switches. The MOTA reached 0.993, 0.991, 0.984, and 0.970 with one to four mice, respectively (**Supplementary methods - Manual validation (extra information); Supplementary Fig. S3b**). As a comparison, a tracking system for pig behavior reached a maximum of 0.90 for the MOTA index^15^.

Finally, it should be noted that beyond this manual validation the system constantly checks the animal identities by constantly comparing the IDs stored by the tracker with those detected by RFID. The system then logs the number of either ID confirmation (match) or ID correction (mismatch) in the database. Therefore, users can monitor the tracking quality of their experiments using the metrics described in supplementary methods or one of their own choice (**Supplementary methods - Automatic tracking control (ATC), ATC based on RFID, ATC based on detection**).

**Figure 1.**
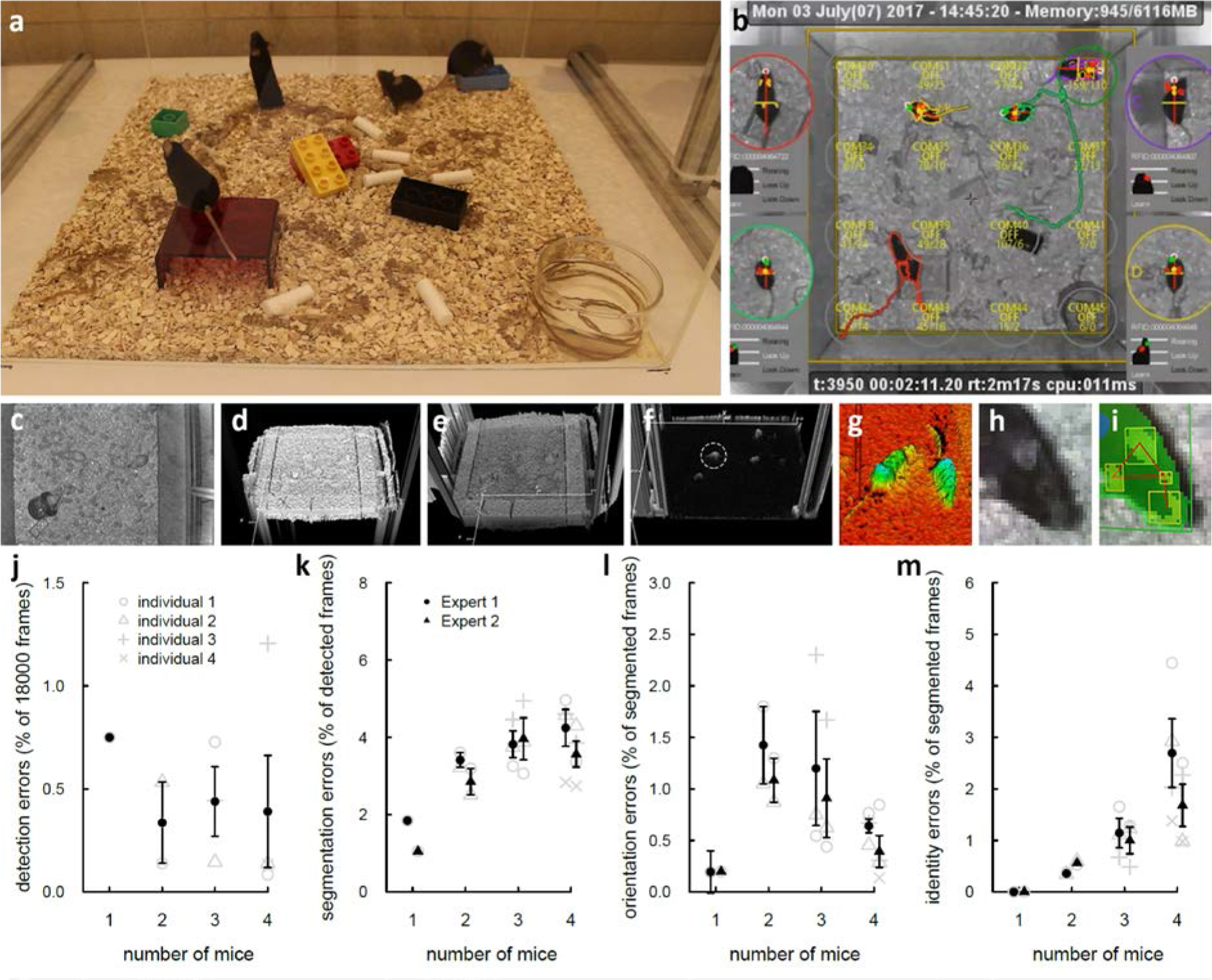
Tracking methodology and validation. **(a)** Real view of the cage with four black mice in enriched environment containing food, water, sawdust, brown crinkle paper, white cotton cylinder, toys, red house (opaque to mice, transparent to infrared). **(b)** Corresponding software display (infrared view). Each mouse is represented with its current segmentation. Colored lines represent tracks during the last 60 frames. Timecodes (experiment and real) are displayed at the top. Circled close-up on both sides represent mice individually, reoriented to head upward. On each close-up, the following information is displayed: the RFID number of the animals, the posture names (rearing, look up, look down; white if active, otherwise black; for instance the yellow mouse is rearing and looking up). The Z-profile of the main axis of the animal with optional detection of ears (red) and nose (green). **(c-i)** Example of segmentation with white animals. **(c-d)** Acquired infrared image and **(e)** corresponding depth map. **(f)** Subtraction of the acquired depth map and background map. **(g)** 3D representation of the detection of the animals. **(h-i)** Close view of the head of the animal with detected nose and ears. **(j-m)** Manual validation performed over 10 minutes with 1 and up to 4 animals. **(j)** Detection error rate: proportion of frames where the animal is not detected. **(k)** Segmentation error rate: proportion of frames where the animal is detected but not correctly segmented (i.e. its shape is not exact). **(l)** Orientation error rate: proportion of frames where the animal is detected and well segmented but head and tail are not correctly detected or reversed. **(m)** Identity error rate: proportion of frames where the animal is correctly detected and segmented but its identity is not correct.

For each timeframe and each detected mouse, LMT provides the mask of the animal, its depth mask relative to the background, the location of the head and of the tail, and finally the ears, eyes and nose detection. This data served as the basis for computing a number of events (list in **Supplementary methods - Behavioral event extraction**), inferred from shape geometry (**Supplementary methods - Event computation**). Overall, we defined 35 behavioral events related to intrinsic and relative positions of the animals and we extracted them automatically. They can be split-up into five categories (**Fig 2a**, list in **Supplementary methods - Behavioral event extraction**). i) Events related to individual behavior reflect activity and individual postures. ii) Social dyadic events reflect the different types of contacts between two mice (see an example of the different types of contacts established between pairs of animals during 4 min in **Supplementary methods - Social contact timelines; Supplementary Fig. S6**). iii) Dynamic events gather the approach, escape and follow behaviors involving two mice. iv) Configuration events reveal subgroup configurations with two, three or four mice, while v) group making and breaking events focus on the dynamics leading to the creation or the ending of subgroups. All these behaviors were computed over the long-term experiments (see example of chronogram in **Fig. 2b** for one individual of a group of four mice recorded for three days). We validated manually the different types of contacts (general contact, nose-to-nose contact, nose-to-anogenital contact, side-by-side contact) since all social events were based on these contacts and on geometric formulae (**Supplementary methods - Manual validation (extra information)**; **Supplementary Fig. S3-5**).

**Figure 2.**
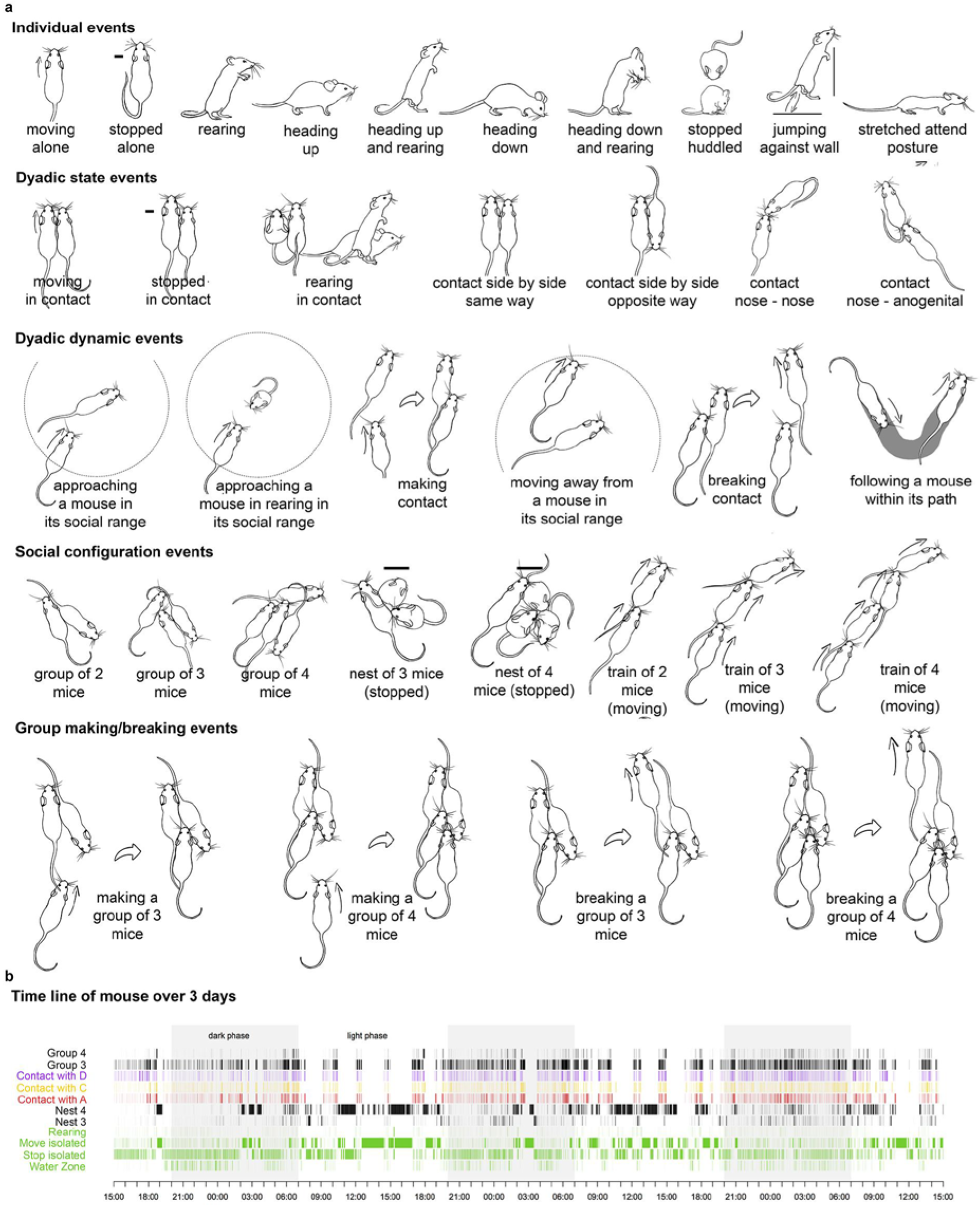
Automatically labeled behaviors and their representation (subset). **(a)** List of behaviors extracted classified in five major groups: individual, social dyadic, dyadic dynamic events, configuration events and group making/breaking events. **(b)** Classification of individual and social behaviors for a *Shank3^−/−^* mouse collected over three days within a group of four mice.

### Individual profiles of *Shank2* & *Shank3* mutant mice

To demonstrate the potential of the system, we compared the behavioral profiles of two mouse models of autism spectrum disorder (ASD) mutated in genes coding respectively for two synaptic scaffolding proteins of the Shank family, namely *Shank2* and *Shank3.* Despite the relatedness of the two proteins, subtle behavioral differences between these two models were expected given the differences in the expression profiles of these two proteins^16^.

We tracked individually four mice within well-established groups to quantify the exploration of the environment and the social interactions within a test cage with fresh bedding and enrichment material. We designed a configuration, based on two wild-type and two mutant mice, allowing a simultaneous monitoring of the control relationship wild-type/wild-type, the interaction wild-type/mutant and the interaction mutant/mutant to provide all possible dyadic relationships that are usually tested with just two animals in classical settings. During 23 hours, we recorded nine mixed-genotype groups of four mice for the *Shank2* strain and during three days six mixed-genotype groups of four mice for the *Shank3* strain. When not specified otherwise in the text, we analyzed only the first 23 hours of the *Shank3* recordings to remain comparable with *Shank2* mice.

In this analysis, LMT automatically extracted 33 behavioral traits for each individual mouse (see list in **Supplementary data - Behavioral event extraction**). These traits were the ones that could be compared between wild-type and mutant mice. The distribution of a subsample of behavioral traits (**Supplementary Fig. S8**) revealed that *Shank2*^−/−^ mice were more affected than *Shank3*^−/−^ mice compared to their respective wild-type littermates. Indeed, *Shank2*^−/−^ mice moved significantly more and spent significantly shorter time in side-by-side contact with individuals of the same genotype in comparison with wild-type mice (**Supplementary Fig. S8**). None of these traits more specifically social events were significantly affected in *Shank3* mutant mice, confirming that *Shank3* mutant mice displayed only subtle behavioral abnormalities^17,18^. To compare mutant and wild-type animals on the same baseline and to avoid inter-experiment variability, the value of each trait (in duration or in number of events) for one mutant was compared with the mean level of this trait over the two wild-type mice of their respective cage. We could therefore detect whether this specific behavioral trait was increased or decreased in mutant mice in comparison with the mean level of wild-type mice tested in exactly the same conditions (i.e., within the same cage).

The behavioral profile of *Shank2*^−/−^ mice (**Fig. 3a**; see **Supplementary Fig. S9** for individual data) reflected higher locomotion activity (time spent moving alone but also in contact, time spent stopped alone), and lower exploration (duration of Stretched Attend Posture - SAP) for individual events. In social events, *Shank2* mutant mice displayed reduced time spent in side-by-side contacts (same and opposite way), slower social approaches leading to a contact (make contact duration), increased following behavior (duration of train2, duration of follow), and increased frequency of completing or breaking a group of three or four mice (make group3, break group3, make group4). In contrast, the behavioral profile of *Shank3*^−/−^ mice (**Fig. 3b**; see **Supplementary Fig. S10** for individual data) indicated significantly reduced activity (moving alone) and significantly reduced time spent in a complex social interaction involving three mice (i.e. time spent at the end of a long line of three mice (train3)). We next compared the behavioral profiles of the two mouse models mutated in genes of the same family to identify the subtle effects of these mutations. Significant differences emerged between the *Shank2* and the *Shank3* profiles in activity measures (Wilcoxon tests with Bonferroni corrections for multiple testing: time spent moving alone [W=213, p<0.00003] and in contact [W=214, p<0.00003], and time spent stopped alone [W=14, p=0.00003]), in exploratory measures (time spent in SAP [W=29, p=0.00044]), in time spent in side by side contact (opposite [W=28, p=0.00036] and same way [W=32, p=0.00079]), in the duration of social approaches (duration of making contact W=190, p=0.00024) and in the dynamic of group3 (making W=200, p<0.00003 and breaking group3 W=196, p=0.00006). Overall, despite carrying mutations in genes from the same family, these two mouse models of ASD displayed inversed phenotypes regarding activity and exploration as well as different alterations of social behaviors.

**Figure 3.**
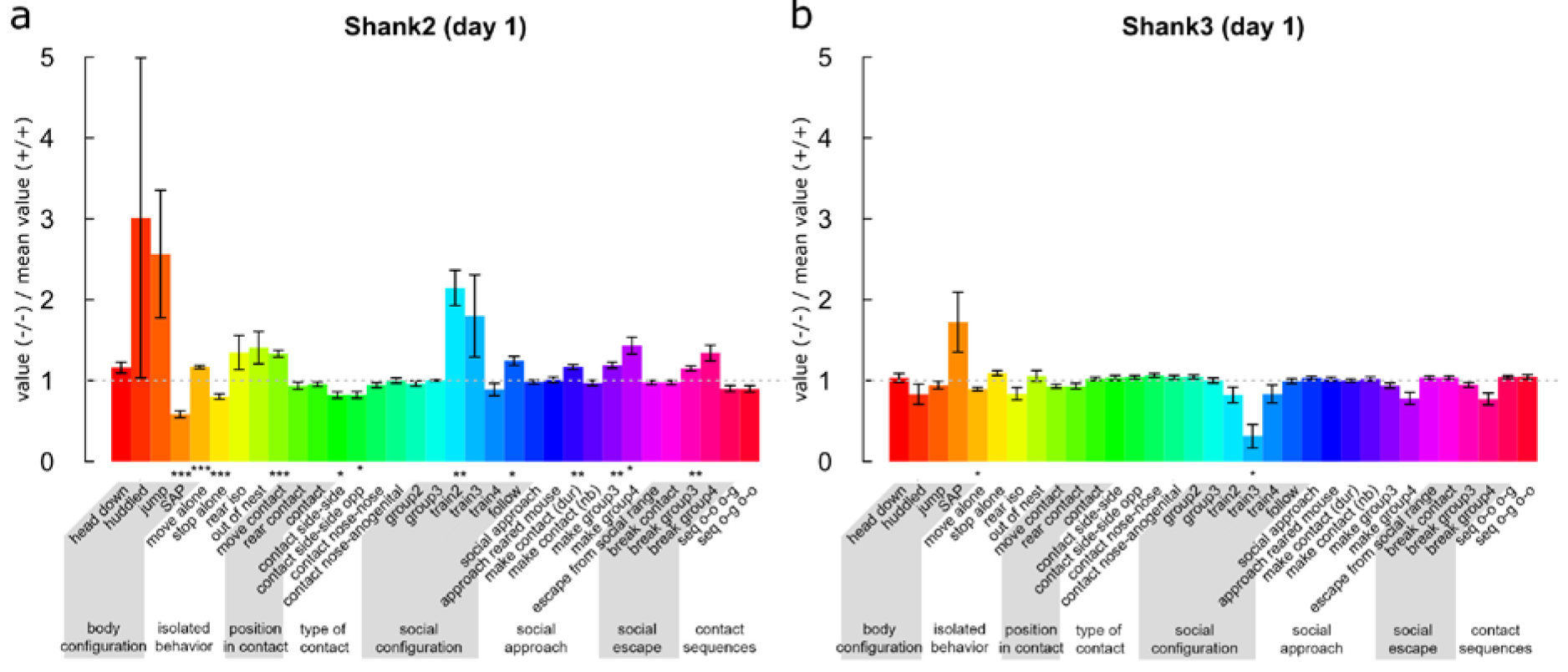
Behavioral profiles of *Shank2*^−/−^ mice (a) and *Shank3*^−/−^ mice (b). Over the first day of group monitoring, 33 behavioral traits were computed for each individual (see definitions in **Supplementary methods - Behavioral event extraction**). Each behavioral trait value for one mutant mouse divided by the mean of the two wild-type mice within each cage was compared to one using Wilcoxon tests (corrected for multiple testing, since 33 tests were conducted for each strain; after correction: *: p<0.05, **: p<0.01, ***: p<0.001). Traits that were not different from the mean value of the wild-type mice of the experiment were set at one. Traits that were expressed more in mutant mice than in the mean of wild-type mice had values larger than one, while traits that were expressed less in mutant mice than in the mean of wild-type mice had values smaller than one. Y-axis graduations represent the expression ratio between mutant and wild-type mice, i.e., a value of 2 represents a trait that was expressed two times more in the mutant mice than in the wild-type mean mouse.

We then investigated in further details specific aspects of the differences between the two mouse models and first focused on specific social behaviors. We conducted these investigations in the same experiments by computing the data extracted from the long-term monitoring of a group of four mice.

### Exploring the subgroup dynamics

In mouse models of ASD, we expect mutant mice to exhibit deficits in social features such as remaining more frequently isolated from the rest of the group in comparison with wild-type mice. We thus provided a comprehensive analysis on the motivation of each mouse to join or leave a particular social structure. Here we define as a social structure the periods when the animals form groups, i.e., when they are in contact or close proximity with one another (see configuration events and group making/breaking events **Fig. 2a**).

We focused on group of three mice, and first noticed that the two types of groups of three mice (two wild-type and one mutant mice or two mutant and one wild-type mice) were equally frequent. This suggested that, contrary to our expectations, *Shank2*^−/−^ or *Shank3*^−/−^ mice were not more frequently isolated than their wild-type littermates when a group of three mice was formed within the cage (**Supplementary Fig. S11a,b**). This also revealed that the hyperactivity of *Shank2*^−/−^ mice did not seem to perturb their ability to form a group of three mice.

**Figure 4.**
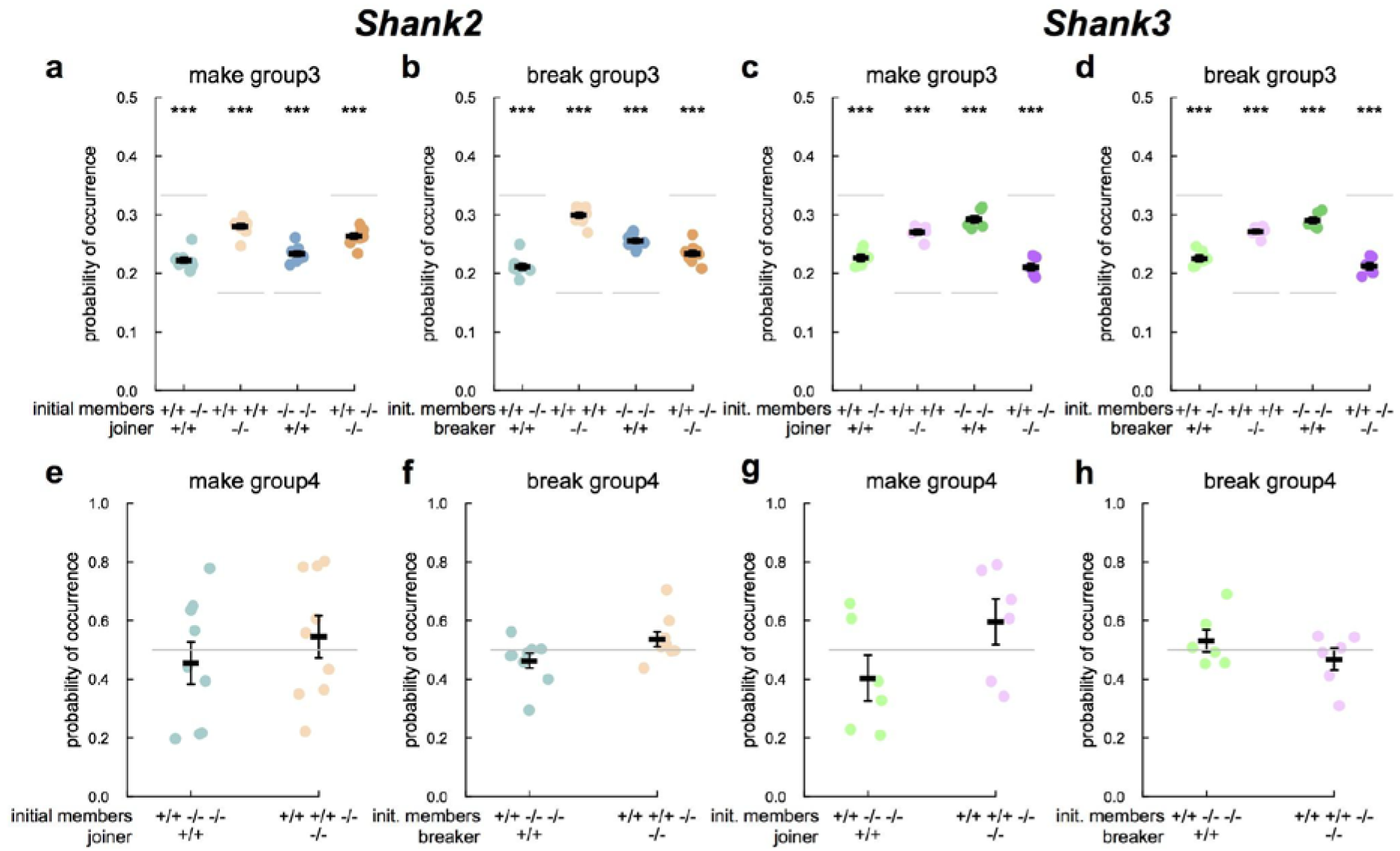
Dynamic of group of three and four mice in *Shank2* and *Shank3* mice. Frequency of occurrence of the completion and breaking of a group of three (upper panels) or four (lower panels) mice according to the genotype of the maker or breaker. Expected probabilities of occurrences are depicted in light grey. With two wild-type (+/+) and two mutant mice (−/−) within the cage, there were 12 possible creations of groups of three mice. For instance, there were two possibilities of a pair of wild-type being joined by a mutant mouse (i.e., 2/12=0.167; Fig. 4a, peach color points). Symmetrically, there were two possibilities of a pair of mutant mice being joined by a wild-type mouse (i.e., 2/12=0.167; Fig. 4a, steelblue color points). There were four possibilities of a pair of one wild-type and one mutant being joined by a wild-type mouse (i.e., 4/12=0.333; Fig. 4a, light blue color points), and four possibilities of a pair of one mutant and one wild-type being joined by a mutant mouse (i.e., 4/12=0.333; Fig. 4a, orange color points). T-tests: *: p<0.05, **: p<0.01, ***: p<0.001.

We then examined group dynamics to assess whether mutant mice were less attracted by social interactions involving more than one mouse. To this end, we first determined in groups of three mice which individual completed or broke it (**Supplementary methods - Study of the dynamics of subgroups of three mice within groups of four mice**). We defined the chance level as the probability of an individual with a given genotype to join or to leave a group of mice over all possible combinations. With two wild-type (+/+) and two mutant mice (−/−) within the cage, there were 12 possible creations of groups of three mice when considering the initial members and the joiner or breaker (i.e. ordered combination, see legend **Fig. 4**). Variations to chance levels indicated that one genotype was more (in the case of higher probability than chance level) or less (in the case of lower probability than chance level) likely than the other genotype to join or to leave a group. In both *Shank2* and *Shank3* strains, the probabilities of a mutant to join/leave a pair of wild-type mice or of a wild-type mouse to join/leave a pair of mutant mice were higher than expected by chance (joining: t-tests after Bonferroni correction for multiple testing: WT-WT<-KO: *Shank2:* t=23.324, df=8, p<0.001, **Fig. 4a**; *Shank3*: t=22.564, df=5, p<0.001, **Fig. 4c**; KO-KO<-WT: *Shank2*: t=14.45, df=8, p<0.001, **Fig. 4a**; *Shank3*: t=19.574, df=5, p<0.001, **Fig. 4c**; breaking: t-tests: WT-WT->KO: *Shank2*: t=28.342, df=8, p<0.001, **Fig. 4b**; *Shank3*: t=28.704, df=5, p<0.001, **Fig. 4d**; KO-KO->WT: *Shank2*: t=21.708, df=8, p<0.001, **Fig. 4b**; *Shank3*: t=23.566, df=5, p<0.001, **Fig. 4d**). Therefore, in both models, pairs of same genotypes (either mutant or wild-type) seem to be more attractive and more repulsive to the other genotype than expected by chance. Since both strains were similar despite their difference in activity levels, this mutant/wild-type distinction could not be explained by hyper-or hypo-activity. Interestingly, in both strains, the mean duration of a group3 event (KO-KO-WT or WT-WT-KO) created by a mouse of a given genotype and broken by a mouse of the same genotype tended to be shorter than group where the joiner and the breaker were of different genotypes (Wilcoxon signed rank tests with Bonferroni correction for multiple testing: *Shank2*: p<0.05; *Shank3*: p<0.1; **Supplementary Fig. S12**). This suggested that group3 created and broken by the same individuals were shorter than those created and broken by different individuals. These short group3 events also reflected the fact that mice have a tendency to just pass near a group of two without stopping.

We also explored the completion and breaking of groups of four animals. The probability to enter as the last or to get out as the first of a group of four mice was not significantly different between mutant and wild-type mice in both strains (**Figure 4e-h**). This suggested that the level of activity did not disturb the social grouping ability of *Shank2* or *Shank3* mutant mice when all mice in the cage are involved.

Altogether, these results indicate that our methodology allows us to access the dynamics of group formation, an information that, due to the technical limitations of current techniques, has rarely been described (except in ^19^). Such possibilities are of high interest for mouse models such as *Shank2* and *Shank3* mutant mice that display a large variability in the severity of their social phenotype according to their genetic construction and to experimental conditions (reviewed for *Shank2* mutant mice in ^20^ and for *Shank3* mutant mice in ^18^).

### Exploring activity levels in groups

The comparison of the individual profiles between *Shank2*^−/−^ mice and *Shank3*^−/−^ mice revealed opposite activity levels (see above), confirming results obtained with classical protocols^8,17^. By following individually the group-housed mice with our system, we were also able to assess how these differences in activity level impact on the circadian activity. Over 23h of monitoring, *Shank2*^−/−^ mice travelled significantly longer distances (5107±300 m) than their wild-type littermates (3121±134 m; Wilcoxon rank sum test: W=12, p<0.001). *Shank2*^−/−^ mice first displayed a novelty-induced hyperactivity at the beginning of the recording. However, despite their strong hyperactivity, *Shank2*^−/−^ mice exhibited typical circadian rhythms, as suggested in another model of *Shank2* mutant mice (deleted in exon 24^21^). Indeed, active and resting periods in *Shank2*^−/−^ and *Shank2*^+/+^ mice were synchronous and the general locomotor activity over 23h was significantly correlated between *Shank2*^+/+^ and *Shank2*^−/−^ mice (Spearman’s rank correlation: rho=0.951, S=2660, p<0.001; **Supplementary Fig. S13**). *Shank3-*^/-^ mice tended to travel shorter distances over the first day (2970±75 m) in comparison with wild-type littermates (3287±116 m; Wilcoxon rank sum test: W=101, p=0.101). The activities of *Shank3*^+/+^ and *Shank3*^−/−^ mice were significantly correlated (Spearman’s rank correlation: rho=0.958, S=2292, p<0.001) and did not suggest that *Shank3*^−/−^ mice were hypoactive specifically because of the novelty of the environment but were constitutively hypoactive. We prolonged the monitoring of these mice over three days to confirm this hypoactivity. Over three days, *Shank3*^−/−^ mice travelled significantly shorter distances (7631 ±142 m) in comparison with *Shank3*^+/+^ mice (8374±226 m; Wilcoxon rank sum test: W=112, p=0.020). Again, the active and resting periods were synchronous between *Shank3*^+/+^ and *Shank3−/−* mice (Spearman’s rank correlation: rho=0.964, S=61024, p<0.001; **Supplementary Fig. S13**). Overall our data revealed a novelty-induced hyperactivity and a general hyperactivity embedded in classical circadian rhythms in *Shank2* ^/-^ mice. This hyperactivity measured by the computation of the total distance travelled confirmed the increased time spent moving (either alone or in contact with any other mouse) and reduced time spent stopped alone identified in the individual profile (**Fig. 3a**). In *Shank3* mutant mice, we observed a general hypoactivity embedded in classical circadian rhythms, which also confirmed the reduced time spent moving alone in the individual profile (**Fig. 3b**). Interestingly, the locomotor hyperactivity of *Shank2*^−/−^ mice might perturb exploratory behavior according to the reduction of SAP behavior (**Fig. 3a**), while locomotor hypoactivity did not seem to have any effect on exploratory behavior (**Fig. 3b**). We therefore next addressed the dynamic of the hyper/hypoactivity in a new environment and the interaction with exploratory behaviors.

### Exploring activity levels and object exploration strategies in single and dyadic tasks

Our system was flexible enough to investigate these differences of activity levels in single (*Shank2* and *Shank3* mice) and dyadic (*Shank2* mice) exploration tasks in detail. During the 30-min habituation in the test cage filled with fresh bedding (phase 1; **Fig. 5a**), *Shank2*^−/−^ mice travelled significantly longer distances in comparison with *Shank2*^+/+^ littermates (Wilcoxon rank sum test: single: W=0, p<0.001; paired: W=0, p<0.001; **Fig. 5b,c**, phase 1). This data corroborates results obtained with the same animals with the classical openfield test (Spearman correlation between the open field test and the single exploration test: rho=0.727, p<0.001; data not shown). The level of activity measured in pairs was significantly correlated with the level of activity measured during single exploration of the test cage (Spearman correlation: phase 1: rho=0.899, p<0.001). In contrast, *Shank3*^−/−^ mice travelled significantly shorter distances in comparison with their wild-type littermates in phase 1 (Wilcoxon rank sum test: single: W=81, p<0.001), confirming their hypoactivity in a novel environment.

**Figure 5.**
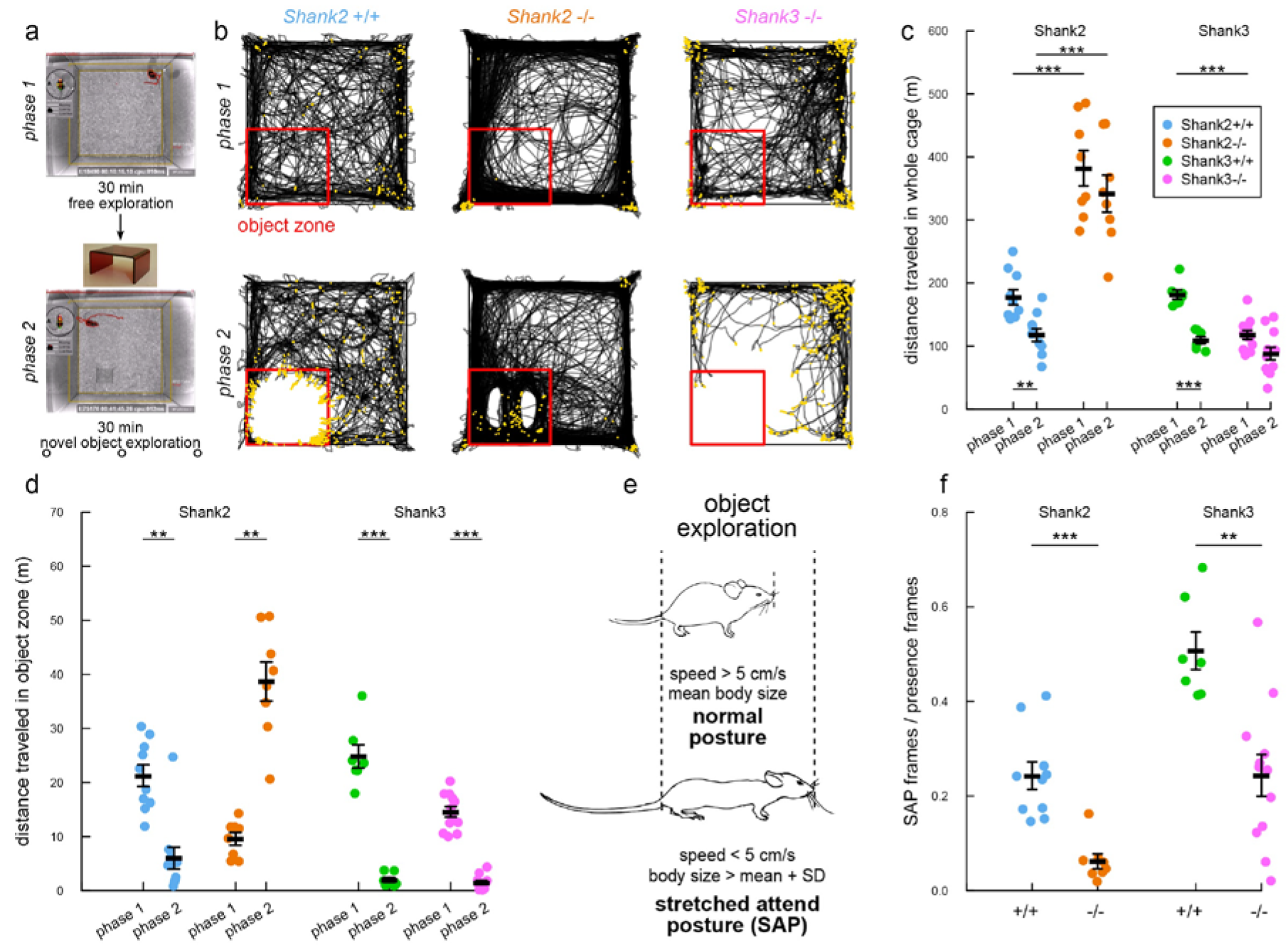
Hyperactivity and atypical exploration strategy in *Shank2*_−/−_ and *Shank3*^−/−^ female mice. (**a**) Protocol used with a single mouse tracked by the system. After 30 min habituation to the test cage (phase 1), a novel object (house in red Plexiglas) was introduced for 30 min (phase 2). (**b**) Examples of trajectories during phase 1 (upper panels) and phase 2 (lower panels) for a wild-type mouse (left panels), a *Shank2*^−/−^ mouse (middle panels) and a *Shank3*^−/−^ mouse (right panels). Red square: object zone; yellow points: frames in which the mouse is in a stretched attend posture (SAP). (c) Distance traveled in the entire cage in phase 1 and phase 2 for wild-type, *Shank2*^−/−^ and *Shank3*^−/−^ mice in the single condition. (d) Distance traveled in the object zone in wild-type mice and *Shank2*^−/−^ mice in phase 1 and phase 2 in the single condition. (e) Definition of the stretched attend posture. (f) Proportion of frames in which mice are in SAP in the object zone over the total number of frames in which animals are detected in the object zone. Data are presented as mean±sem and individual points for 10-12 *Shank2*^+/+^ mice and 8-12 *Shank2*^−/−^ mice as well as 7 *Shank3*^+I+^ mice and 12 *Shank3*^−/−^ mice (Wilcoxon rank sum test; **: p<0.01; ***: p<0.001).

We finally investigated the influence of the activity level on object exploration. For this purpose, after phase 1, we introduced on the fly (i.e. without stopping experiment) a novel object (here a red house) and the mouse could explore it for 30 min (phase 2; **Fig. 5a**). During this second 30-min phase, hyperactivity was still significant in *Shank2*^−/−^ mice in comparison with their wild-type littermates (Wilcoxon rank sum test: single: W=0, p<0.001; paired: W=0, p<0.001; **Fig. 5b,c**). Again, the level of activity measured in pairs was significantly correlated with the level of activity measured during single exploration of the test cage (Spearman correlation: phase 2: rho=0.895, p<0.001). Interestingly, once the object was introduced, *Shank2*^−/−^ mice displayed higher activity in the object zone in phase 2 than in the same zone in phase 1 (Wilcoxon signed rank test: single: V=0, p=0.008; paired: V=0, p<0.001; **Fig. 5b,d**). In contrast, *Shank2+^/+^* mice displayed a decreased activity in the object zone (Wilcoxon signed rank test: single: V=55, p=0.002; paired: V=77, p=0.001; **Fig. 5b,d**), similar to *Shank3*^+/+^ (Wilcoxon rank sum test: W=49, p<0.001) and *Shank3*^−/−^ mice (W=144, p<0.001). This suggests two different exploration strategies: cautious approaches of the object in *Shank2*^+/+^, *Shank3*^+/+^ and *Shank3*^−/−^ mice and uninhibited exploration in *Shank2*^−/−^ mice (**Fig. 5b**, phase 2, lower panels). In paired experiments with the *Shank2* strain, the occurrence of each strategy depended on the genotype of the mouse and was not influenced by the genotype of the mouse they were paired with (see **Supplementary Fig. S14a-c**).

The two different strategies were characterized by the occurrence of stretched attend postures (SAP) (**Fig. 5e**, yellow points on **Fig. 5b**). SAP is a risk-assessment posture^22^ automatically quantified in our system, no matter whether the animals are monitored in single or social conditions. It is characterized by an elongated body (body length longer than the mean body length + one standard deviation) and a reduced speed (< 5 cm/s). In the object zone, *Shank2*^−/−^ mice used significantly less frequently the SAP to explore the novel object in comparison with their *Shank2*^+/+^ littermates (Wilcoxon rank sum test: single: W=78, p<0.001; paired: W=128, p=0.001; **Fig. 5f**), suggesting that *Shank2*^−/−^ mice lacked risk assessment and therefore displayed atypical exploration. The presence of a conspecific did not seem to modulate this abnormality according to the results obtained in the paired condition. We replicated these findings about hyperactivity and atypical exploration strategy in a second cohort of *Shank2* mice including males and females for single object exploration and females only for paired object exploration (see **Supplementary Fig. S14d-f**). This difference in novel object exploration suggests that *Shank2* mutant mice present a suppressed neophobia (i.e. absence of inhibition), but this might be independent from their initial increased anxiety in dark-light test^8^. Surprisingly, *Shank3*^−/−^ mice also displayed less frequently the SAP in the object zone in comparison with their wild-type littermates (Wilcoxon rank sum test: W=77, p=0.002). This suggests that, despite a similar distance travelled around the object, *Shank3*^−/−^ mice still displayed subtle abnormalities in their exploration strategies.

## DISCUSSION

LMT makes it possible to phenotype animals in groups and over long periods of time (days to weeks). This new methodology, based on the acquisition of rich sets of data, opens new avenues for examining how different genotypes, pharmacological tests or enriched environments influence decision-making or social interaction in a robust manner. This system also provides new information on social interactions and more specifically on interactions involving more than two freely-moving mice. Gathering data on several animals simultaneously and over long periods of time generates large datasets highly representative of individual behaviors. This in turn will stimulate massive analysis of large datasets to comprehensively study complex behaviors and should allow a statistical analysis of persistent individual traits. To boost this large-scale approach, LMT provides a website to allow the community to share data and analysis scripts, following the example of MouseTube^23^ (**Supplementary methods - Collaborative website sharing data; Supplementary Fig. S7**).

LMT is a real-time process that also opens new perspectives to interfere online with any behavior by triggering an external device when a pre-determined event is detected. Indeed, the tracker is able to provide, in real time, the location and the posture information of each mouse. We made this data available by low-latency network connection (**Supplementary methods - UDP live network information stream**) so that any third-party device (Arduino-like devices enabling automation control) or third-party software can gather current tracking information, either on the computer performing the tracking or on a dedicated one on a local network. We demonstrate this feature with an example: a live rendering of the subjective view of each mouse in 3D created with unreal engine (**Supplementary movie - LMT - Live 3D rendering demo**). In this toy demo (using only x,y,theta for each mouse), one can see the scene from the point of view of each mouse. We do not, however, claim an accurate reconstruction of the visual field of the mice. In this demo, mice leave a trail of the color corresponding to their identities to display their past trajectories to better understand the inter-individual coordination of movements. This approach can be extended to record electrophysiological activities, ultrasonic vocalizations, physiological signals such as cardiac activity, and also to build closed-loop systems to react to the behavior of the mice at very specific moments with optogenetics or other stimulating systems triggered with the closed-loop signals. The system should therefore be used to develop new behavioral tests to better answer the phenotyping needs^24^.

Finally, LMT provides a built-in repertoire of behavioral events that can be analyzed with scripts as illustrated in this paper. This repertoire includes individual events such as stop or rearing, but also social interaction events such as the different types of contacts or social configuration involving more than two mice. An exhaustive set of additional data, including the positions (x,y,z) of the head, tail and mass center for each animal at each time point, as well as the complete mask of each animal are available for custom extensive and elaborate analyses. This rich description of individual mice opens the possibility to conduct complex offline analyzes on large datasets, such as the analysis of event behavioral sequences^25^, individual movement tracking or social network analysis. For example, among the *Shank2* one-day group recordings that we conducted as pilot experiments, we observed a group of four animals that were not nesting in the provided house, while all other groups nested there. We investigated this case (**Supplementary movie - Investigating an abnormal nest building behavior**). One mouse appeared to avoid the house and refused to get in it. This animal spent most of its time at the opposite location of the house. The other mice first set the nest in the house, and then, after a few hours, moved its location and all the nesting material to the location of the other animal. Finally, the structure of the database complies also with the addition of extra measurements to monitor the environment for further developments. This should increase the reproducibility of experiments, and be used to optimize housing conditions by analyzing behavioral markers of welfare (social isolation, stress) and abnormal behavior.

The low price of the system enables users to multiplex setups and conduct all experiments at the same time, which is an advantage when running long-term recordings. It is worth noting that the total number of animals that can be tracked simultaneously is only limited by the computing power (each individual tracking has a related CPU cost) and the density of mice present in the field (animals need to be alone from time to time to be identified via RFID). In the future, we plan to connect several setups to provide larger environments to larger groups of mice. We encourage the community to take over LMT and we facilitate ways to improve performances. Indeed, thanks to the open-hardware and software framework, and all the blueprints provided on the web, anyone can reproduce the system, build over it and improve or adapt parts of it without having to painfully rebuild everything from scratch. LMT has been designed natively for parallel architectures and to make the most out of new and more powerful computing architectures such as the Ryzen
CPUs. This scale-up in computing power benefits directly the performances of machine learning instances, and improves the identification latency. It will also impact the efficacy of data storage and processing during long-term experiments.

## ACKNOWLEDGMENTS

Yannick Archambeau, Patrick Ollivon at the workshop of the Institut Pasteur for building the 12 first setups and advising on hardware. William Meiniel for mathematical proof in head/tail probability decision. Microsoft France for their technical support. Piernicola Spinicelli for optical engineering and critical reading of the paper. Raphaël Marée for machine learning support. Barbara König for advices and critical reading of biological experiments. Jacqueline N. Crawley for careful reading and comments on the manuscript. Angelos Barmpoutis for providing us the early kinect 2 driver and his very helpful support. Nicolas Chenouard that drove use to a Machine Learning solution. Pascale Dugast for drawing the mice in the different behavioral events. Aaron Engelberg for careful checking of English spelling. Sébastien Wagner and Robert Accardi for RFID advices. Marcio Marim for the website development. Xavier Montagutelli and Marion Bérard for animal facility support. This work was partially funded by the Institut Pasteur, the Bettencourt-Schueller Foundation, the Agence Nationale de la Recherche through grants ANR-10-LABX-62-IBEID and France-BioImaging infrastructure ANR-10-INBS-04, the Centre National de la Recherche Scientifique, the University Paris Diderot, the BioPsy Labex, the Institut National du Cancer, the Foundation for Medical Research (FRM, Equipe DEQ20130326488), the Innovative Medicines Initiative Joint Undertaking under grant agreement no. 115300, resources of which are composed of financial contribution from the European Union’s Seventh Framework Program (FP7/2007-2013) and EFPIA companies’ in kind contribution. The funders had no role in study design, data collection and analysis, decision to publish, or preparation of the manuscript.

## AUTHOR CONTRIBUTIONS

F.d.C. created the tracking and analysis methods, developed the open-hardware setup and software. E.E. performed and analyzed experiments. N.T. analyzed experiments. S.D. provided real-time programming advices. T.La. provided the probabilistic framework for machine learning decision making. T.Le. provided electronic support. A.I. created the blueprints. A.-M.L.S performed the genotyping of the mice. F.d.C., E.E., N.T., P.F., T.B. and J.-C.O.-M. conceived the project and wrote the manuscript.

## DATA AVAILABILITY

Full datasets (databases and films) generated during and/or analyzed during the current study are available in the Live Mouse Tracker website repository, https://livemousetracker.org/.

## CODE AVAILABILITY

Full source code is available at http://icy.bioimageanalysis.org/plugins/livemousetracker. It includes Java code, R analysis Scripts and Python analysis scripts. This also includes CAD hardware resource files.

## SUPPLEMENTARY METHODS

**Supplementary Figure S1.**
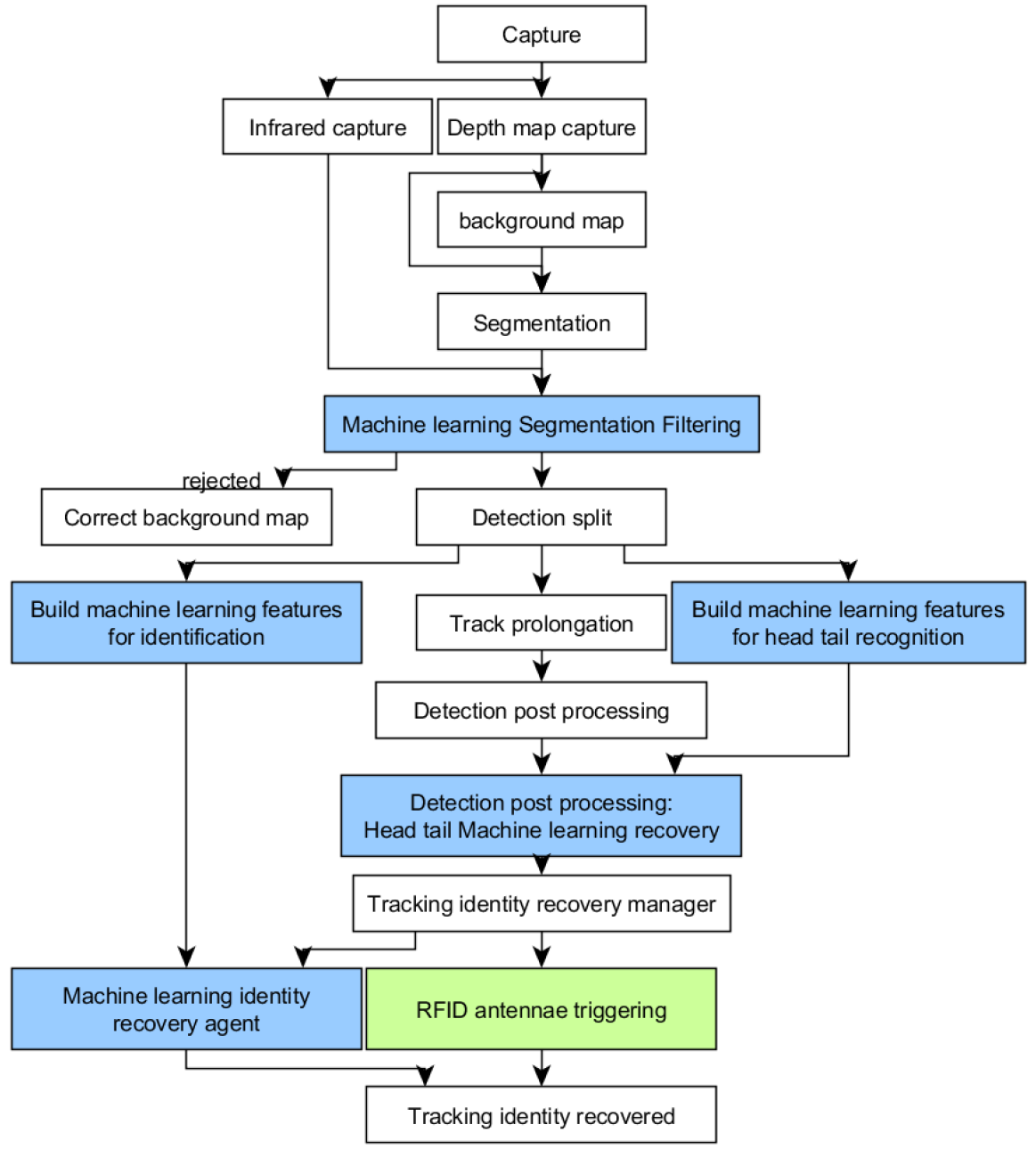
Tracking diagram

### Capturing depth-map

The depth map is represented as an image of 512×424 (typed signed short). We correct invalid values grabbed from the sensor that appear as spikes in the depth map. If the value is within an unexpected range (equals 0 for saturation or less than −32768), we repeat the value of the previous valid pixel read. *Code: LiveMouseTracker.correctInvalidZValue.*

The accuracy of the depth-map measurement is affected by the light absorption of the observed material. The brighter the object is, the farther it artificially appears. The correction is mild (less than 5 millimeters range), but mandatory to observe small animals. We tested 8 different Kinects and found a common empirical linear offset correction using the infrared images. We correct depth-map by applying the following formula to each pixel:

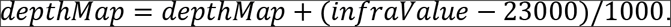

*Code: LiveMouseTracker. compensateZIntensityError.*

The Kinect firmware is originally designed to watch player’s large movements standing in a room (like dancing), and is therefore not designed to observe close objects. Its original minimum firmware working range is set to 50 centimeters, but practical observation below 80 centimeters is not possible. To obtain a closer range, and a better resolution, we mask with a tape one of the infrared blaster of the Kinect to reduce scene illumination. (**See Supplementary movie and file - Blueprint and assembly instructions**). This could also be obtained with optical density placed in front of the blasters.

### Computing the background height-map

The background depth map is represented as an image of 512×424 (unsigned short). For each new depth-map captured by the sensor, we update the background map such that for each pixel, the background map keeps the min value between the captured and the stored maps. Area containing detections are not processed.

*code: livemousetracker.BackgroundHeighMapBuilder.integrateNewDepthImage.*

In case of false detection (see below), the height-map is corrected using the detection mask. For all pixels belonging to this mask, we set the height-map values with the values of the detection mask. Thus, the false detection will no longer be detected.

*Code: BackgroundHeighMapBuilder.correctBackground.*

### Segmentation and detection

The segmentation map represents all the animals and objects detected in the field. We compute the boolean segmentation map as follow: we set the segmentation map as *true* if *heightMap* − *depthMap* > *depthSensitivity*. In our setup, *depthSensitivity* is constant and equal to 14, i.e. 1.4 millimeters. If infrared value is saturated, depth is not reliable and we discard the corresponding pixels from segmentation. We then crop the segmentation mask with the defined region of interest. To obtain all individual segmentations, we perform a connected component extraction (Icy internal *BooleanMask* function). Segmentations are then classified in two lists: spurious and validated detections. Segmentations with less than 30 pixels get in the spurious list (they correspond to sawdust moving in the cage). Segmentations in spurious lists larger than 3 pixels are sent to the *backgroundMapBuilder* to correct the background. Their count is stored for further analysis in the “frame.particle” field in the database as it reflects the sawdust spread by mice when moving, digging or fighting. If the user sets the option “Manage wired animal”, extra filtering is applied, based on ellipse fitting of objects. The longest size of a mouse being around 60 pixels, objects greater than 70 pixels in their main fitted axis are rejected. Then we process the validated segmentation list. If a detection is bigger than *MaxSizeDetection,* it might be because several animals are in contact, therefore the detection is processed by the detection splitter (see below).

Code: *detection.MouseDetector.detectMice()*.

### Machine learning - building the detection feature vector

The feature vector for machine learning is computed for each detection. The feature vector is key as it allows the machine learning to classify objects (instances). Therefore the feature vector must reflect the object and act as its signature; it must also contain enough information to discriminate objects. As a constraint, we tried to find the smallest number of features to have both a good reliability on the signatures and to make the training of the machine learning as fast as possible. Mice do have a large variety of conformations. Roughly, they can be elongated, completely retracted and appearing ball-like, rearing (so only the head is visible), or over an object (appearing bigger). Therefore, we do not want to consider surface, shape descriptor or scale in the learning. We also do not consider direction of the animal nor its location. Note that min/max intensity or depth are also not coded to avoid size or shadow discrimination by the machine learning. We only normalize histograms to make them consistent with or without shadows. The feature vector is composed of 33 values. The first value is the ID of the animal (i.e. its class available only when training machine learning, under supervision). The next 16 values are the infrared histogram and the last 16 values the depth histogram. Code: *machinelearning.MachineLearningSetBuilder.*

Each histogram is built on either infrared data or depth data values corresponding to the mask of the animal. We set 16 bins starting from the min value to the max value of the dataset. Histogram values are then normalized.

*Code:detection.MouseDetection.buildHistogram.*

### Detection filtering with machine learning

The segmentation process detects all the objects that are moving and only filters very small objects. However, other objects are moving in the cage: food, house, large area of sawdust when animals dig, enrichment objects during object discovery tasks. Therefore we need an additional filtering able to determine whether we observe objects or animals. This filtering is performed by a supervised machine learning (random forest (1000 trees, depth 100), *weka* library http://www.cs.waikato.ac.nz/ml/weka/). The machine learning is trained using the first seconds of the observation, where moving objects are expected to be part of the class “Animal” versus a random pick of patches labeled as “Others”. The maximum number of observations considered is 600 per class, which represents 20 seconds of observation of a single animal or 5 seconds of observation of 4 animals as they are all mixed in this classification. This training set is then updated continuously in a background process using the last detection of each animal. Detection filtering is performed for each detection at each frame using a predictor of the machine learning. If *p*(*detection is class animal*) > 0.7 the detection is set as “Animal”, else as “Others”. If we still have more items detected than the maximum number of animals expected, we remove the smallest detections. For each detection, a test is performed to find the number of tracks which terminates close to the detection. If several tracks match a single detection, it means that two or more animals are getting in contact, so we need to split the detection before the tracking association process starts.

### Detection splitter

The detection splitter tries to recover animals that are melted in a single segmentation. It uses tracking information to get the number of tracks that ends at the melted segmentation. Then the splitter uses the last detection found at those locations and depth-map data to split the detection in the number of animals expected (i.e. the number of tracks ending at the segmentation). To perform the split, we first create an index map, mapped on the segmentation mask that will store the identity index of each pixel in the mask. The index map is initialized with the pixels representing the main axis of each previous detection. If the main axis is not available, we initialize with the mass center. Then we process the dilatation of the index following two constraints within the same process. The first one is a 3D constraint: each dilation is performed z-slide per z-slide, starting from the higher altitude of the segmentation down to the floor. Second, at each z-step, dilatation processes constrain the number of pixel dilated to maintain an equal share of all the available pixels between animals so that final splitted segmentations tend to have the same size.

Code: *package livemousetracker.splitter.*

### Tracking extender association process

Once detection set is complete, we first try to prolong existing tracks. We get all detections at *t* − 1. If the detection is less than 30 pixels away, we extend the track. If several tracks are sharing potential prolongations, we perform an hungarian algorithm to set the best assignment. If no track is found, we create a new track.

Code: *livemousetracker.track.TrackExtender.*

### Identity recovery with machine learning

The tracking identity recovery process is a multi-agent process performed as a background task. The tasks addressing the tracks to identify are scheduled by a manager process working with several solvers (Identifiers) in parallel. Depending on the number of threads available on the computer, we affect a given number of identifiers to the manager. The manager will dedicate one identifier to solve identities of unknown tracks in the present, and all the remaining identifiers will be launched to solve identities of anonymous tracks randomly located in the past. A manager builds the identifiers and affects them the anonymous tracks. It also guarantees that identifiers are not working on overlapping tracks. Code: *livemousetracker.identity.MultiIdentityAgentManager.*

An identifier agent receives the track to identify and retrieves concurrent tracks, identified or not, that are overlapping in time with it. The identifier attempts to provide the identity of all the tracks involved at once, even if several (or all) tracks are anonymous. For each track involved, the identifier creates a “track identity scorer” (TIS). For reliability purposes, TIS dealing with tracks shorter than 2 seconds are discarded. Then, for each track involved in the TIS, we create a set of possible identities that is initialized with all identities available. This set of solution is pruned by removing all the identities found in overlapping tracks. Then the TIS trains a dedicated machine learning restricted to the animal remaining in its solution set. This machine learning is trained with a random pick of 5400 detections from each animal (equivalent to 3 minutes of observation per animal). We then pick 60 detections from the track managed by the TIS, and for each detection we use the machine learning predictor which gives a vector representing for each detection its probability to belong to each animal. Within the next steps, all those results will be multiplied together. Nevertheless, due to a lack of training, the machine learning sometimes does not recognize a detection at all if it is an observation outlier and subsequently gives a probability of 0. In that case, this value would set to a probability of 0 a whole association hypothesis. Thus, to avoid this kill switch, we arbitrary floor this probability result to 0.01. To speed up the TIS process, we cache (i.e. store) the custom machine learning created by each TIS so that it can be directly queried by other TIS without re-training it. Meanwhile, to adapt to any change of appearance of the animals or to new conformation displayed, caches are destroyed 2 minutes after they have been created.

The last step is called the Track Identity Global Solver (TIGS). We create recursively all the possible tracking identity association of all the tracks with every pruned animals per track. Detections scores of all tracks per animal are gathered from all the Track Identity Scorers. For each possible global solution, it provides a vector containing the 60 * *number of animals* detection evaluation. We then compute the product of the whole vector. Once we gather all the detection results for all association, we compute its sum, and the ratio of the best association towards that sum. We take the association decision if the final association ratio is greater than 0.95. In that case, we apply the solution found and affect identities to all tracks. If the condition is not satisfied, the identity is not recovered and anonymous track are left anonymous. The Identity Manager will therefore start later on a new Identifier on those tracks. Meanwhile, the knowledge of the machine learning will be updated with new observations that may better fit the data that remain to be identified, meaning that future observations can solve past tracks. Note that for memory consideration, data are streamed to database after 5 minutes, meaning that if no identity is found during 5 minutes, track will be saved as anonymous, flushed from memory and will never be identified.

Code: *identity.TrackIdentityGlobalSolver.*

### Head/tail detection post-processing

Mice are mostly moving forward. They can appear to go backward when they are rearing, during jump phases along walls or when they are running over a wheel. To automatically find the head location of the mice, we fit an ellipse over the animal and get its main vector. One of its extremities is the location of the head. The animal mask is then cut into two sub-masks (A and B). We therefore need to find whether A or B corresponds to the head part of the animal. We use the speed of the animal to set the head mask, therefore we need to know in which track this detection is associated to compute its speed. Thus, head/tail is processed in post track-processing.

Code: *MouseDection.testA_B_MC_SpeedForHeadLocation()*.

To perform this recognition, we train one machine learning per animal (in a background task) using the same features computation as for the detection. We keep 300 rolling detections to feed the machine learning, but we enable its query as soon as we have 150 detections (5 s of observation). As each machine learning is dedicated to an animal, they can look completely different, or be equipped with different devices on their head and body. Once the head/tail machine learning of the animal is ready, we use the predictor to compute *pAA* = *p*(*A is ahead*) and *pBB* = *p*(*B is back of the animal*). If *pAA* * *pBB* < 1 − *cst* and 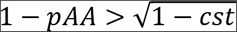 and 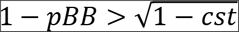 with *cst* = 0.85, we perform the swap.

Code: *mousedetection.PostProcess().*

### Head sub-parts processing

The head sub-parts processing is only possible if the mouse is not wired. The head subparts are detected depending on the fur color of the animal. We first detect if the animal is white or black. Given that the signal provided by the kinect is very stable (from one experiment to another, in time, and also from a kinect to another kinect), and that the contrasts between the parts of an animal are important, we can use hard-thresholding. We extract the mean intensity of the shape of the animal in the infrared image. If the score is below 8000, we set the mouse as black, otherwise it will be white. If the animal is black we threshold the mask in the infrared image at 4000. This will detect spot corresponding to ears, nose and eye (and potentially legs). We then check all detections in order to test if a subset satisfies distance constraints. The distance between the ears should be in the interval [7,16] and symmetrical, and the nose should be at the extremity of the animal. For white mice, we detect only the eyes by thresholding the infrared image (25000). To determine gaze direction, we need ear and nose extraction, therefore it can work only with black mice. To detect the tilt of the head, we project the ear/nose location on the main vector of the mouse. Then we compute the ratio of the vector tail to ear/nose projection. If the ears ratio is over 0.9, it means that they are located close to the front edge of the animal, therefore the animal is looking down. If the nose is detected, we set the animal as “looking up”. All other configurations lead to no labelling of neither “looking up” or “looking down”.

Code: *mousedetection. MouseDetection()*.

### Camera setup and calibration

The Kinect camera should be facing the floor. Its front edge should be at 63 centimeters from the floor (floor is bottom of the cage considering no sawdust) to get the best resolution for a 50×50 cm cage. The Kinect should be connected to an USB3 port. The Kinect should also be taped to enable close observation as detailed in supplementary blueprints. The
*LiveMouseTrackerCalibration* program (demonstrated in the supplementary movie *Hardware blueprint and assembly instruction*) helps by positioning the camera perpendicular to the floor (**Supplementary Fig. S2**). It provides a live matrix display of distance measurements to finetune the pan/tilt of the camera. It also warns if the sensor is not taped. The provided setup provides images at a resolution of 1 pixel = 0.175 cm for an object observed at 63 centimeters from the Kinect.

**Supplementary figure S2.**
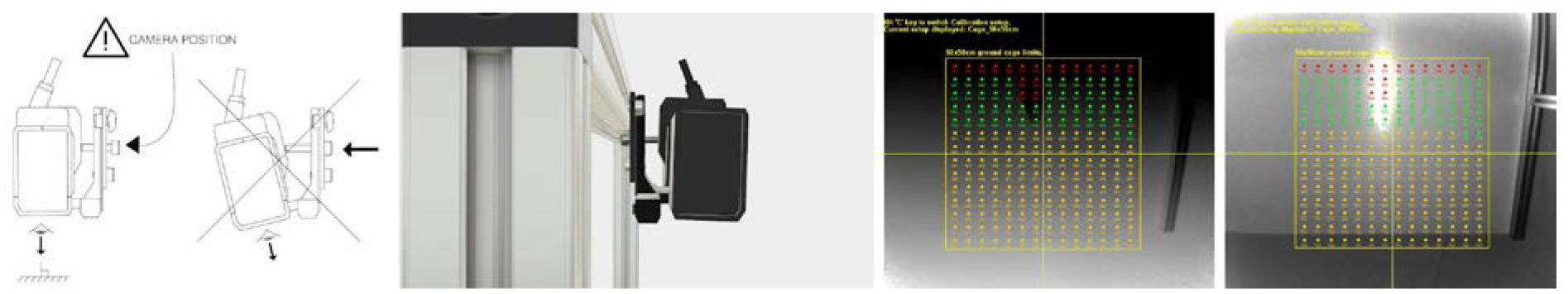
Calibration of the setup. (a) The notice from the assembly instruction which indicates that Kinect should be perpendicular to the ground. (b) 3D image from blueprints. (c-d) View of the calibration software. All colored dots display live the distance to the ground so that the user has just to move the Kinect by hand to watch if it is properly positioned. Red, orange, and green dots indicates respectively very bad, bad and correct positionnement.

Note: The kinect v2 is now discontinued. Millions of items have been sold so this is not a rare product. Nevertheless, the method can be applied with other RGBD cameras if the driver to connect to the Java core program is available.

### RFID calibration

The reading frequency of the antennas should be at 134 kHz, or 125 kHz depending on the RFID probed used (the frequency used is region dependent). Frequency should be accurate at −1/1 kHz to get the best reading range. This study and the provided blueprint have been designed and performed with 134 kHz probes (Glass probe ISO 11784/11785 2×12mm). Antenna reader enables the self measurement of the antenna reading frequency in a process called “Measure Unit Operating Frequency” (MOF). Use the provided *RFIDReaderTest* program while soldering to tune its frequency by adjusting wire length. Extra capacitor should be soldered on the reading board to switch to 125 kHz reading.

### Triggered-RFID closed-loop control

To read a passive RFID probe carried by an animal, the antenna induces an electric current in the probe which powers it. Then the probe modulates the received signal to transfer its identification back to the antenna. This whole process is performed in 100 milliseconds by the RFID reader hardware (code at *rfid.RFIDAntenna.run*). If several probes are powered up at the same time (i.e. several animals in the range of an antenna), the signal is jammed and no ID is read. The signal is also jammed if several antennas closer to 70 centimeters are activated at the same time. The RFID identification is in closed-loop with the video tracking. The software data structure contains a list of antennas with their respective COM port number, position and range in image field. It activates in priority antennas to query identities of anonymous tracks. We obtain all detections available at current *t* from the anonymous trackpool. We remove the animals that are in contact in this list as the reader will be jammed if two RFID are present over it at the same time. Then we activate the antenna that is the closest from a detection. Once a reading is performed, we correct the existing tracks and solve possible track conflicts (i.e. several tracks having the same ID at one timepoint). Machine learning instances operating in the background to solve identity of tracks are then canceled. If no antenna is activated with the previous process (meaning all animals are identified or they are all in contact), we use the same procedure to read identities of known animals to validate constantly their identity. code: *rfid.RFIDManager.activateAntennas.* If an antenna gets faulty during the process, the user can see it on the interface (the corresponding antenna is drawn red). An antenna can get faulty because of an inopportun USB unplug or loss of power. The program is communicating with each antenna within its own process to avoid any lock while transmitting data. If a timeout is reached, the antenna is set as faulty. To monitor the quality of each antenna, the interface displays the number of reading attempts and the number of effective probe readings for each antenna.

### RFID hardware

RFID can be bought at http://www.priority1design.com.au ref FDX-B/HDX RFID Reader Writer with external antenna and USB port (30€). Coil for antenna is ref RFIDCOIL-100A. (3.5€). The system is best working with 16 antennas/readers distributed on the 50×50 cm area. See Supplementary Blueprint and assembly instructions.

### Database description

All the data are stored locally on a simple standalone database, in a redundant manner to target different levels of query, which complies with the Big Data principles^26^. This means that the database can be accessed by any third party software such as R, Matlab, Python or Java and that the user can query the database on simple or complex features. No server is needed to store or query databases as the data are shared between experimenters by simple file transfer (contrary to dedicated database servers that would lock-in the data into labs).

The software creates the database for the current experiment automatically as the tracking starts. We choose the SQLite database format as it is server free. The whole database is a single file, and each database represents an experiment, making it easy to backup, share, query and transform for users. Tables are Animal(Id, RFID, Genotype, Name), RfidEvent(Id, RFID, Time, X, Y), Frame(FrameNumber, TimeStamp, NumParticle, Paused), Event(Id, Name, Description, StartFrame, EndFrame, IdAnimalA, IdAnimalB), Detection(Id, FrameNumber, AnimalID, Mass_X, Mass_Y, Mass_Z, Front_X, Front_Y, Front_Z, Back_X, Back_Y, Back_Z, Rearing, Lool_Up, Look_Down, Data). The data field is an XML text containing the full mask of the animal in zip format. To perform fast SQL Queries, we advise to add indexes to the database (Provided in **Supplementary SQL script**). Database can be explored with the software DB Browser for SQLite (http://sqlitebrowser.org/). We also provide scripts and examples to create all the figures and query shown in this paper from the software R (https://www.r-project.org/). Code: *experiment. Experiment. createDataBase.*

### Manual validation (extra information)

In the following figure (**Supplementary Fig. S3**), we detail the distribution of the mean duration of error and the representation of the MOTA index^14^ depending on the expert.

**Supplementary Fig. S3.**
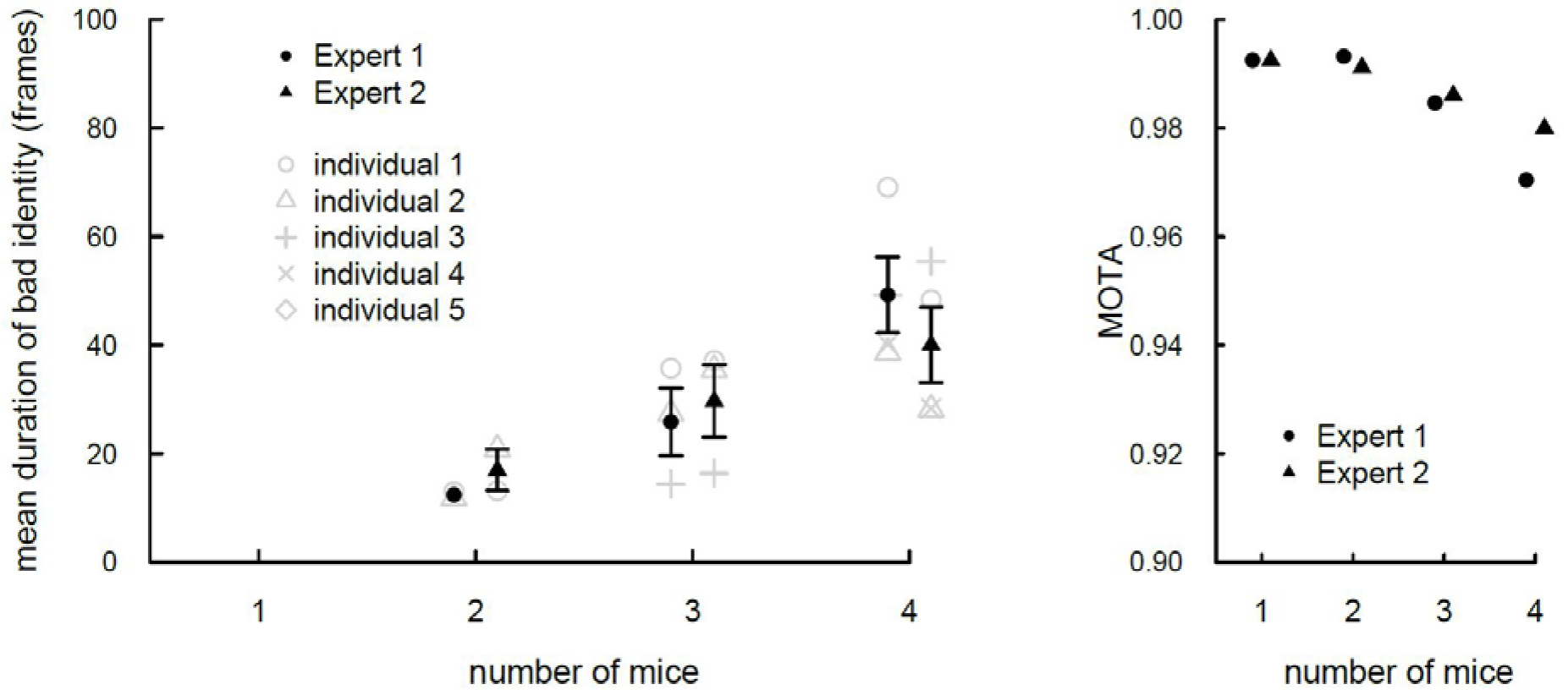
Manual validation of the tracking system. (a) Mean number of consecutive frames where the identity of a mouse is not correct. (b) Calculation of Multiple Object Tracking Accuracy (MOTA): MOTA = 1 - sum(t) ( false negative(t) + false positive(t) + identity switch(t)) / sum(t) (nb mice in the ground truth), where t is a frame, a false negative is a frame where a mouse is not detected but it is there, a false positive is a frame where a mouse is detected but it is not there, an identity switch is a frame where identities are switched between individuals, and the number of mice in the ground truth is the number of mice that should be tracked in each frame.

**Supplementary Fig. S4.**
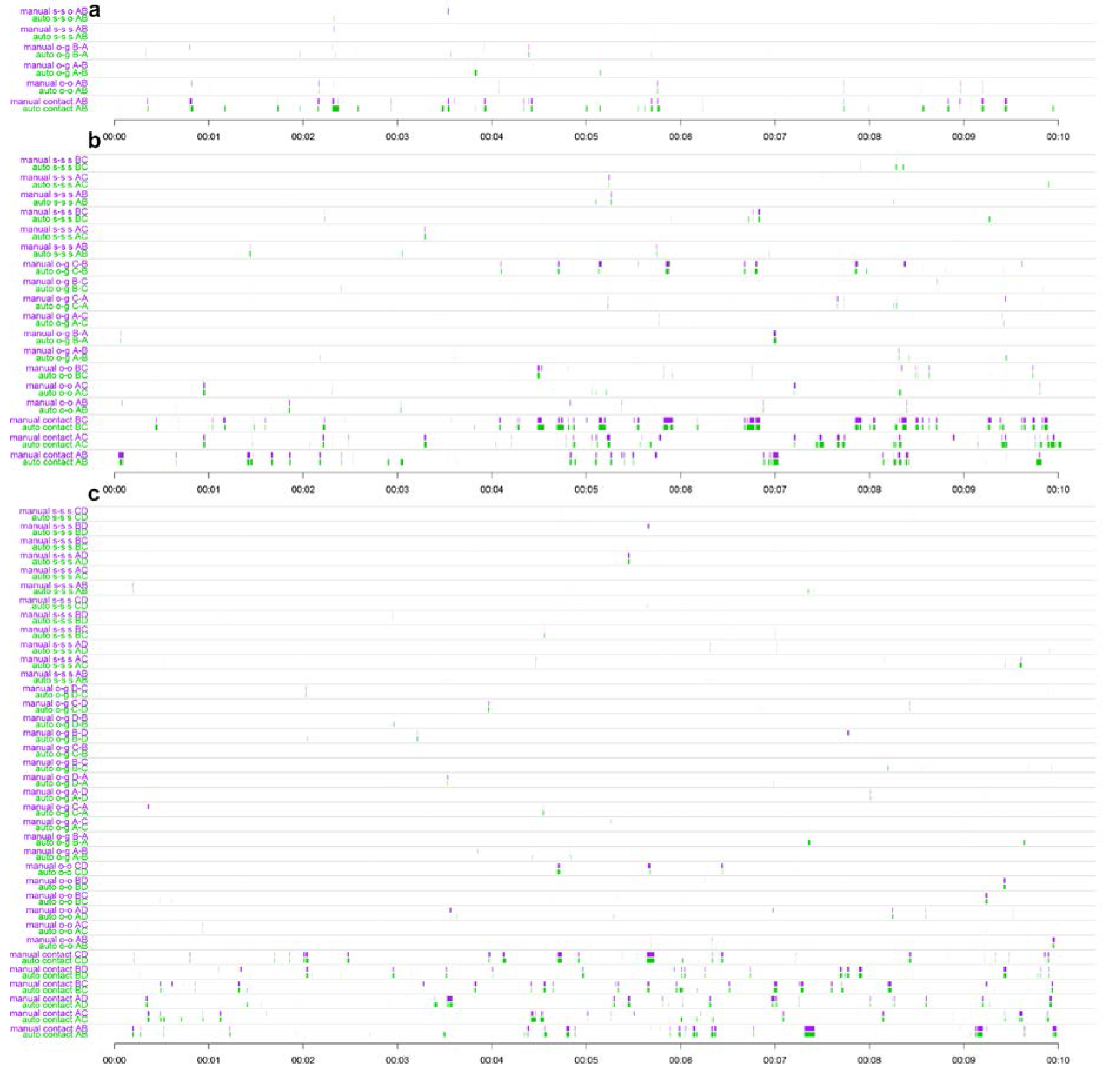
Timelines of manual validation of the tracking system (10 minutes). Each consecutive couple of row displays the manual validation (violet) against the same behavior labeled automatically (green) over 10 minutes. (a) Experiment with 2 individuals. (b) 3 individuals. (c) 4 individuals.

**Supplementary Fig. S5.**
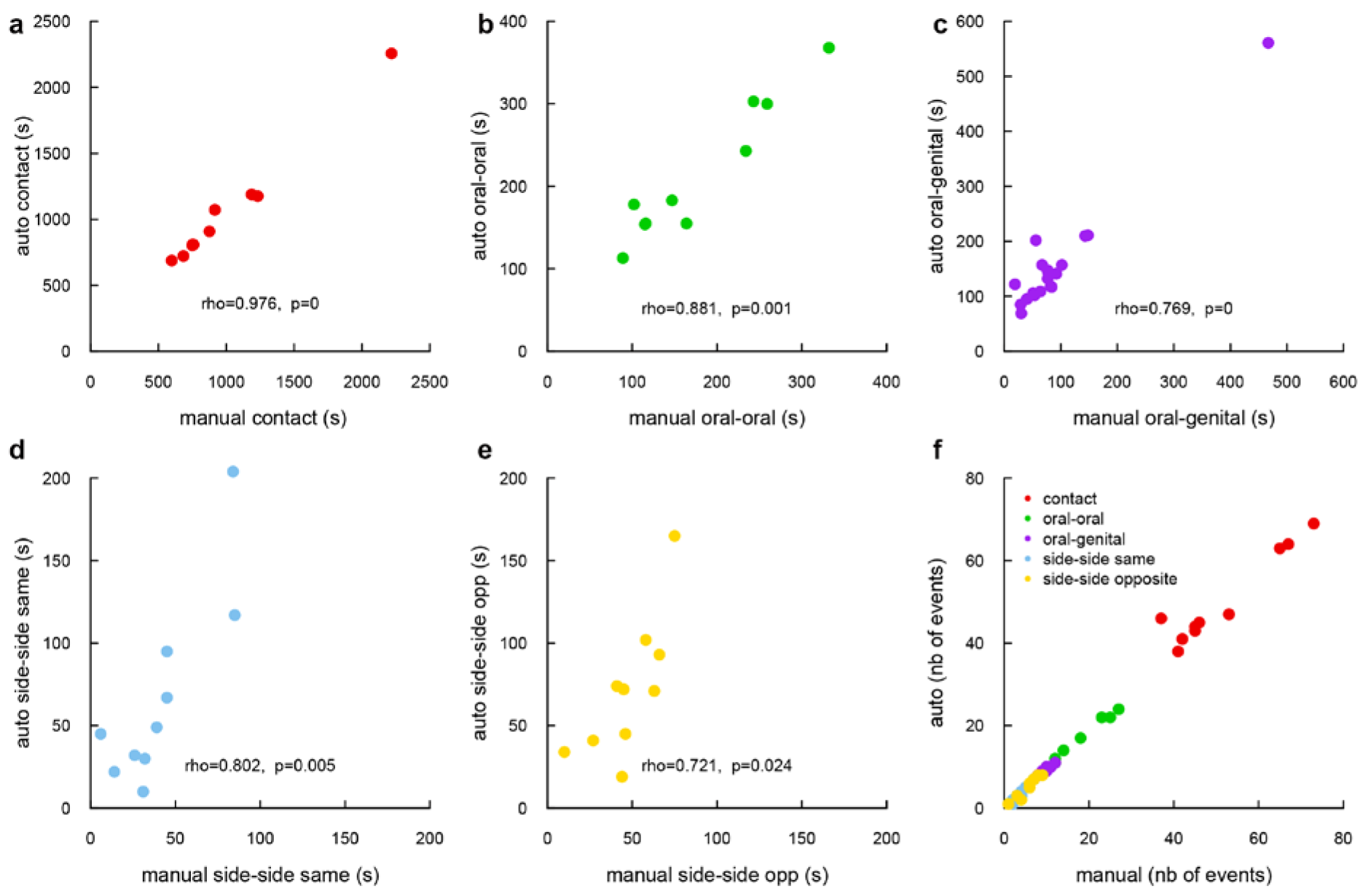
Manual validation of the tracking system, (a-e) Correlation between automatic and manual measurements for the different types of contact: (a) general contact, (b) oral-oral contact, (c) oral-genital contact, (d) side by side contact in the same direction and (e) side by side contact in the opposite direction (Spearman’s rank correlation). (f) Number of events that were detected automatically as a function of the number of manual events detected by the experimenter.

The **Supplementary Fig. S5** displays the correlation (in total length of event found) between the manual and the automatic labeling on five different contact events. The **Supplementary Fig. S5a-e** displays linear correlation. For the time spent in contact, the time spent in oral-oral contact and the time spent in oral-genital contact, the manual and the automatic results display a significant positive correlation and therefore match. For the side-by-side contact events, the automatic method tends to find longer events than the manual method, but the relation between the automatic measure and manual measure is still linear. After further inspection, we realized that our expert was a little more restrictive on its criteria for side-by-side contact events, which explains the reduced duration of manually measured events in comparison with automatic measures. We checked this assumption by comparing the two measures at the level of the event. For each manually labeled event, we searched for an event in the automatic labelisation. The **Supplementary Fig. S5f** shows that very few events were missed by the automatic labelisation, which confirms our explanation.

### Automatic tracking quality control

Live Mouse Tracker records events in the database of the experiment to monitor the tracking. Five types of events are stored: 1. If the machine learning dedicated to the identification of tracks performs an association, an event “MACHINE LEARNING ASSOCIATION” is logged. 2. For the RFID, we log three events. a) If an anonymous track is assigned thanks to RFID, then “RFID ASSIGN ANONYMOUS TRACK” is triggered. b) If a RFID read corrects the identity of a track, then the “RFID MISMATCH” is triggered (and the track is corrected). c) Finally when all animals are identified, the RFID are used to control the identity of animals. In that case, at each reading confirming current track identity, a “RFID MATCH” is stored in database. 3. We can also monitor if an animal is not detected at a given frame. This happens when animals are hiding, or when they become indistinguishable, for instance in the nest condition. In this case, the event “DETECTED” is not triggered for the given animal. Those events (built automatically) allow building of several measures (online or in post-processing) to assess the quality of the tracking. Here we provide two different measurements:

### Automatic tracking quality control based on RFID reading

We extensively uses the RFID technology to assess the reliability of the identity. In our experiments, the identity of each animal is checked through the RFID antennas 49 times per minute on average (from 19.7 to 100.9) (see supplementary table 1). At each RFID read, the identity of the animal is checked and corrected if necessary (the correction propagates back in time). In the table hereafter, the number of identity match over the total number of reads provides **98.47% ± 0.16 (mean ± SEM)** of success. The following table details the score for all animal of all 23h-experiments (**Supplementary Table 1**).

**Supplementary table 1.**
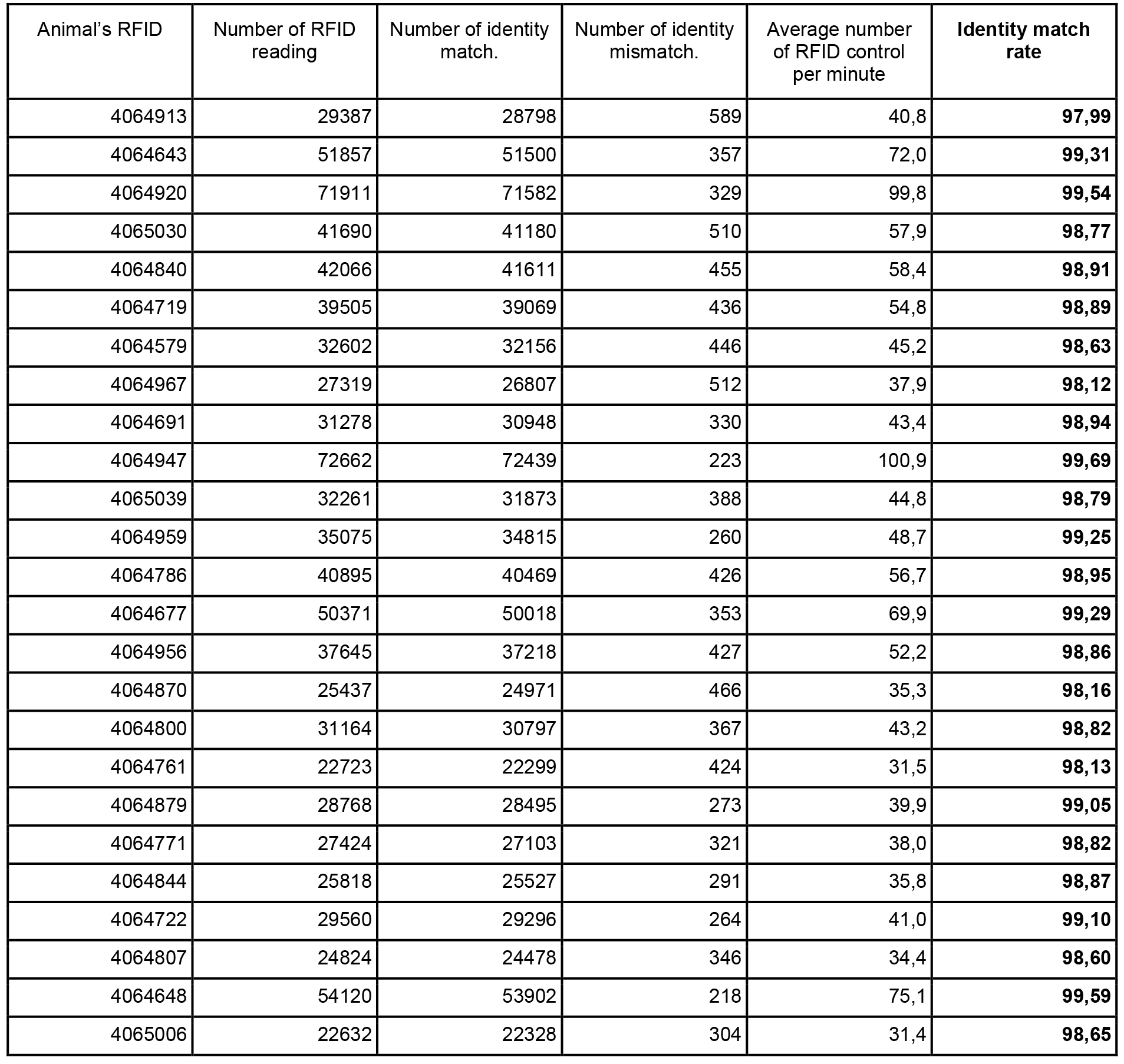
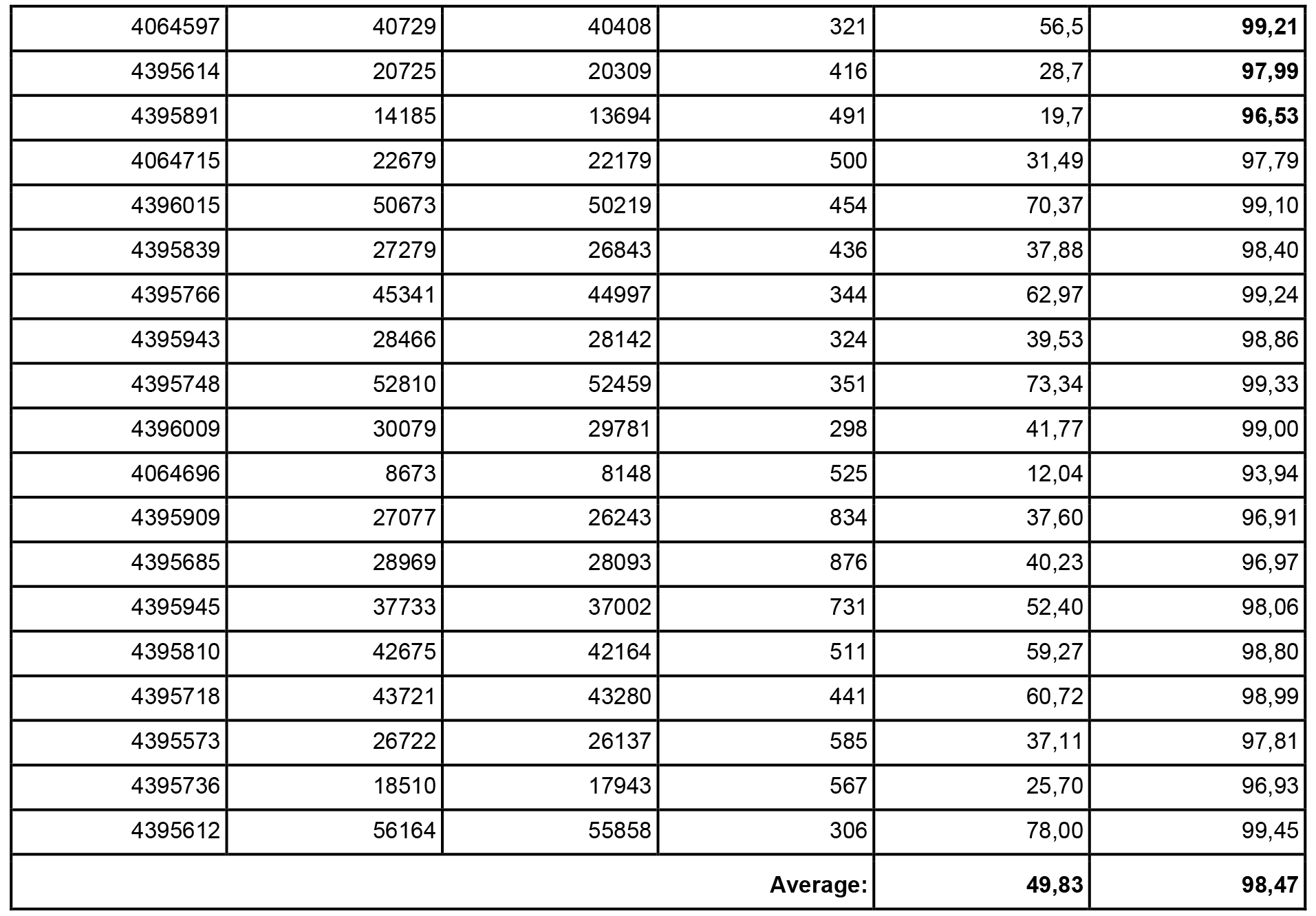
RFID check quality for 12H (all Shank2 mice)

### Automatic tracking quality control based on detection

The second quality metric considers the number of frames the animal is detected over the total number of frames of the experiment (as used in validation) (**Supplementary Table 2**). In our experiment, we observe animals in housing conditions. They cannot escape the observation. So we observe different group activity that can be condensed in 2 phases: activity or sleep. During sleep, the animals are all together in a nest for long periods, and may be covered by cotton. We assume that our system cannot track individuals during those phases. Still, we detect those events and we label such periods as “nesting”.

Therefore the quality measurement during activity provides **92.91% ± 0.48 (mean ± SEM)** of success.

**Supplementary Table 2.**
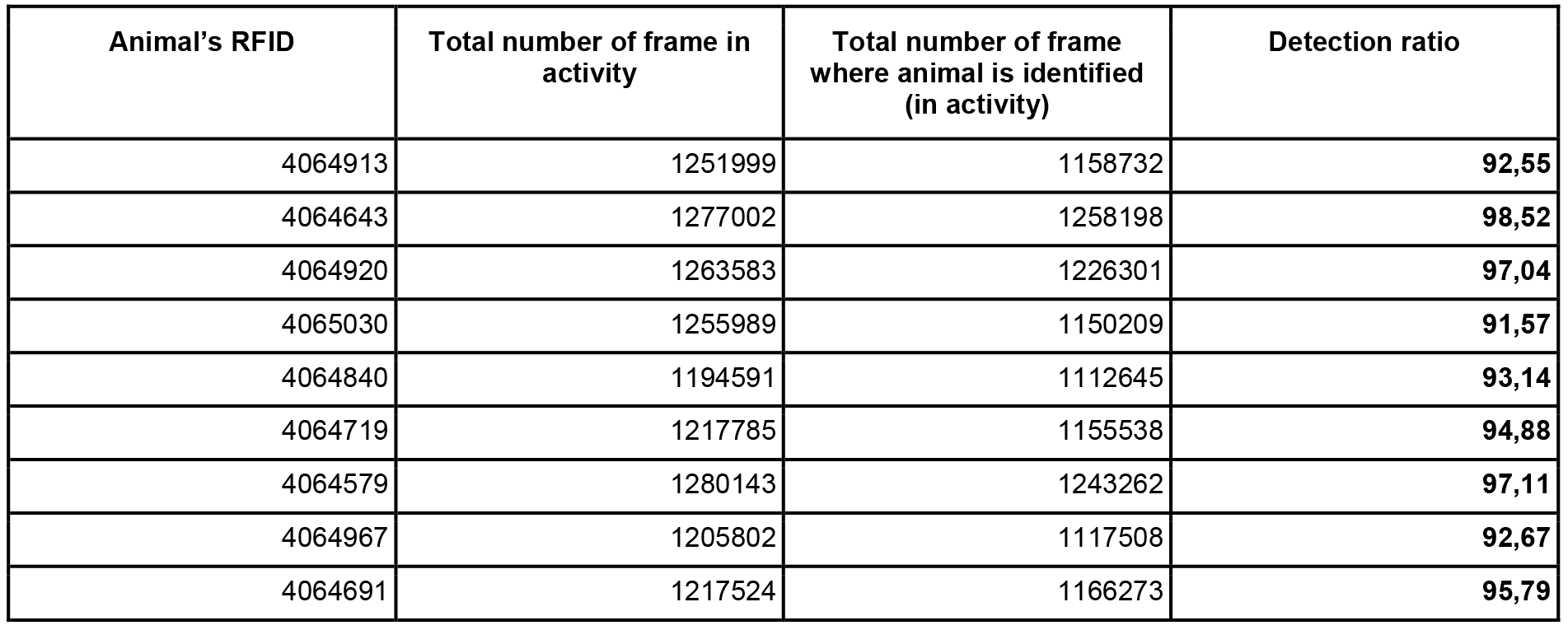
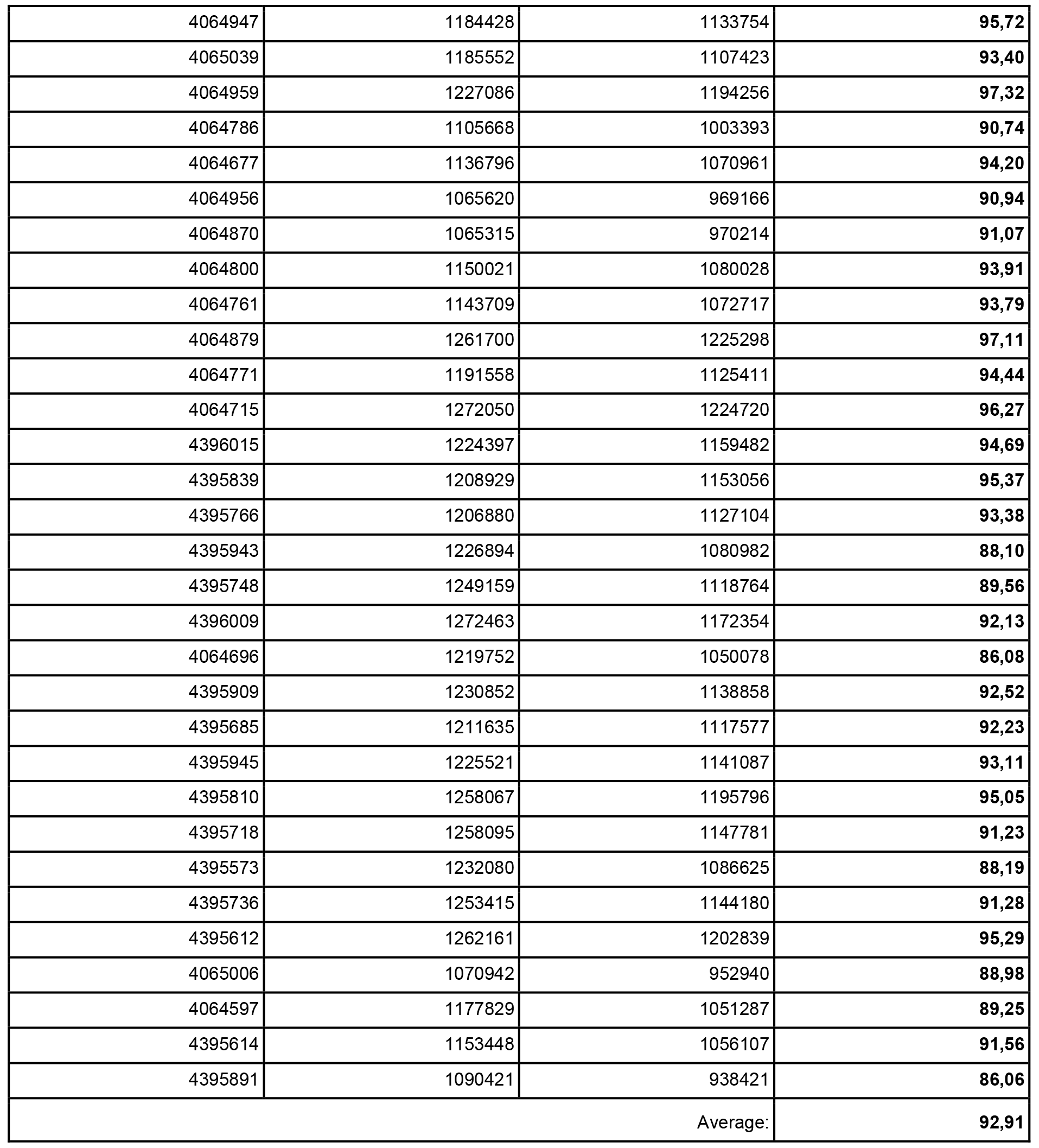
Detection check quality for 12H (all Shank2 mice)

### Social contact timelines

In the following figure (**Supplementary Fig. S6**), we provide additional timelines for one experiment of 4 individuals.

**Supplementary Fig. S6.**
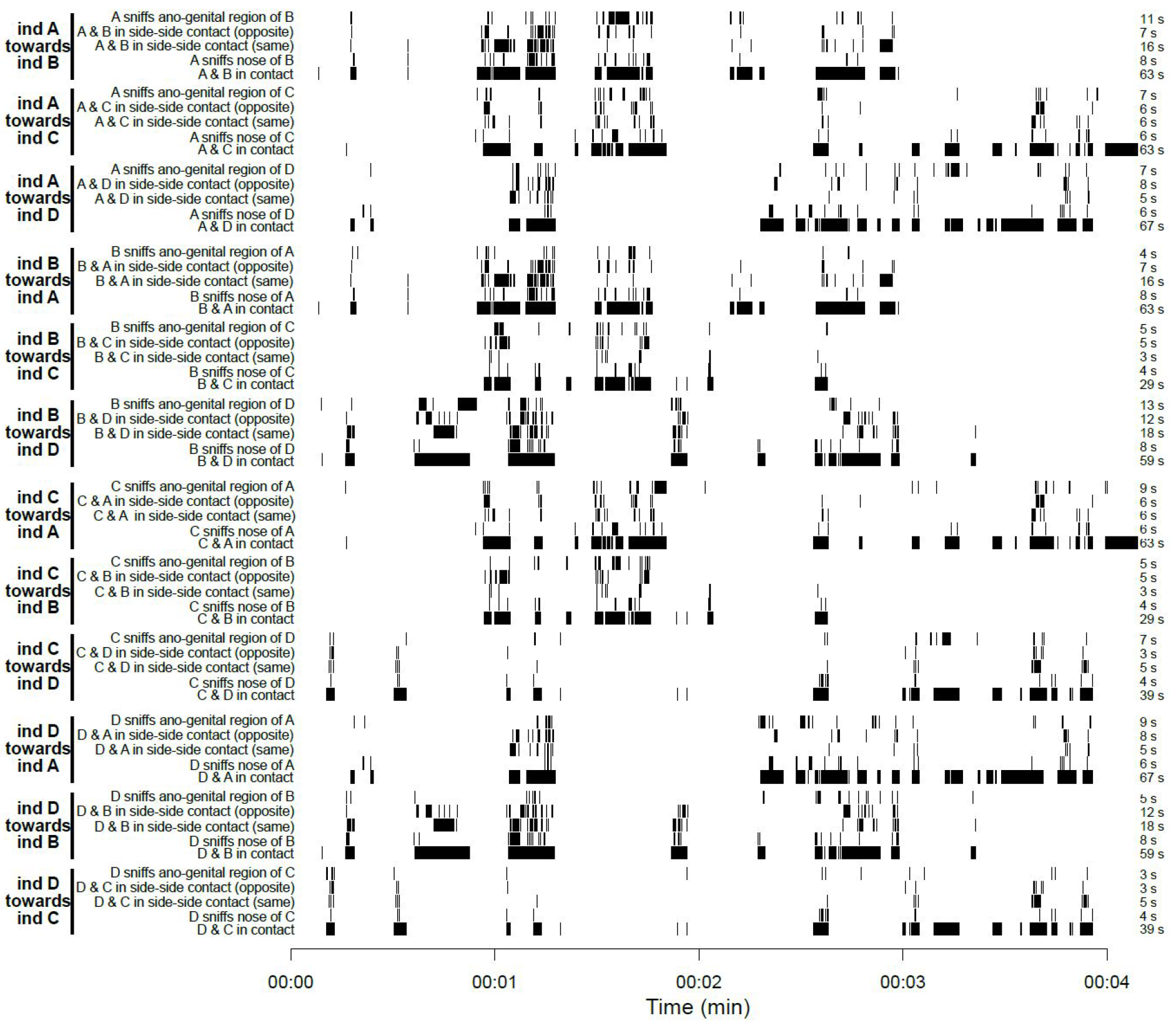
Example of the distribution of the different types of contacts between each possible pair of individuals within a group of four mice.

### Querying database information with R and Python

Supplementary files are provided. They consist in scripts displaying how to query databases with R and Python. We also provide examples of scripts to build new events.

### UDP live network information stream

To perform closed-loop control over any device, we deliver live tracking information, streamed on network. To avoid network packet control, we choose the UDP protocol, which is efficient for continuous data streaming. Streaming message consists in location and direction of all animals present in the field.

### Behavioral event extraction

We extracted individual, dyadic, configuration, dynamic as well as making / breaking group events (see examples in **Fig. 2**). The following list describes the events that were calculated. Events preceded by a star were used for the computation of the individual profiles.

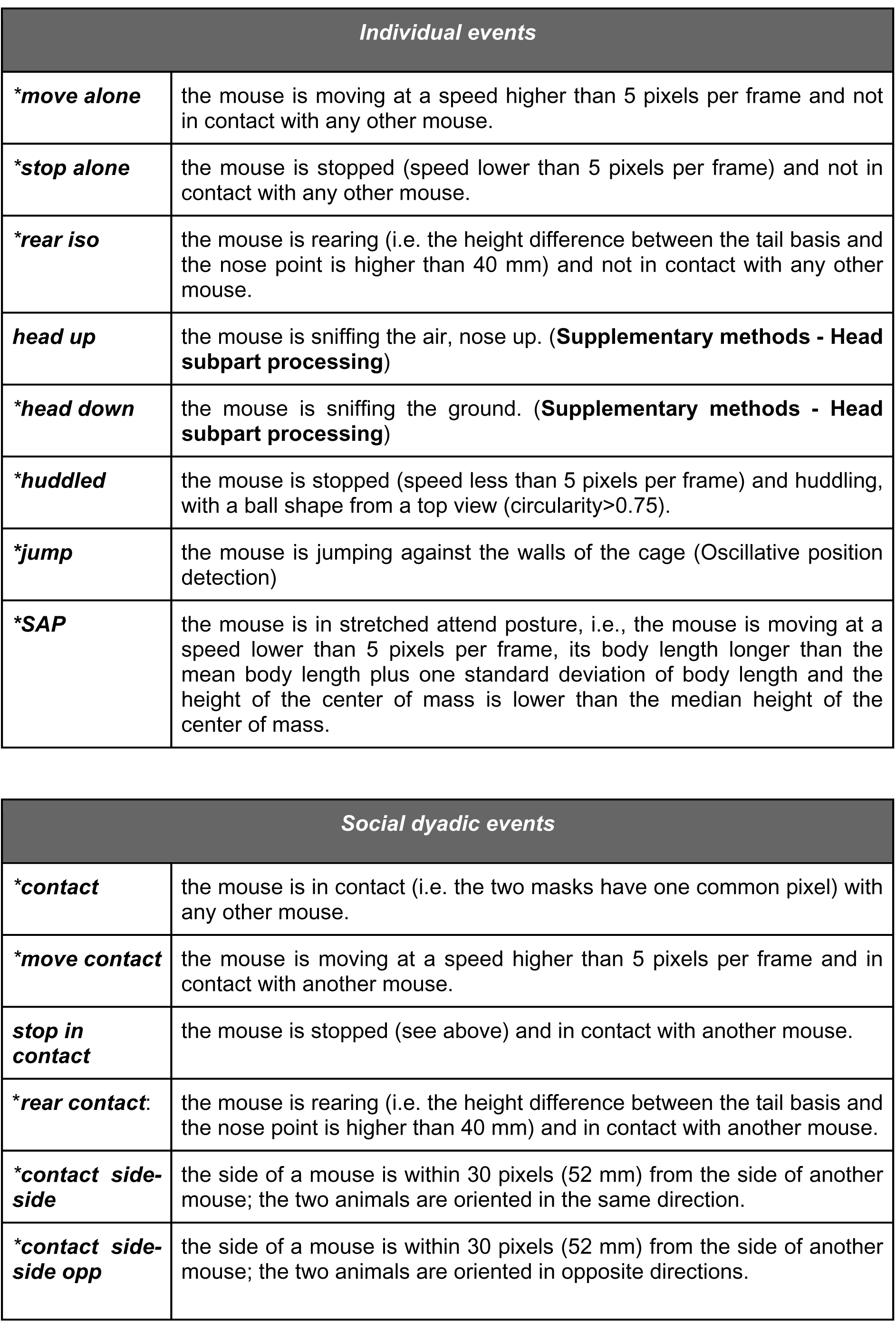

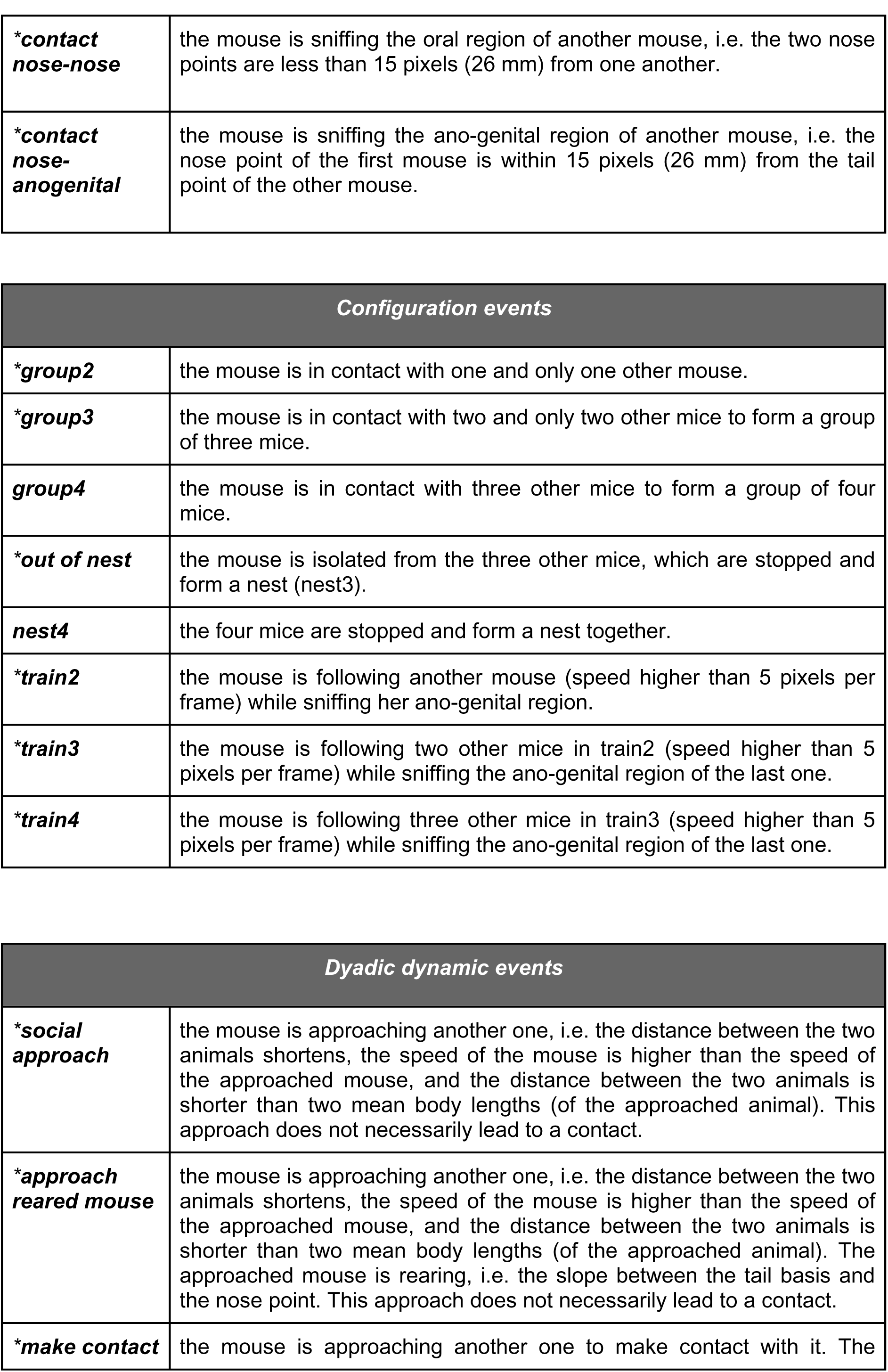

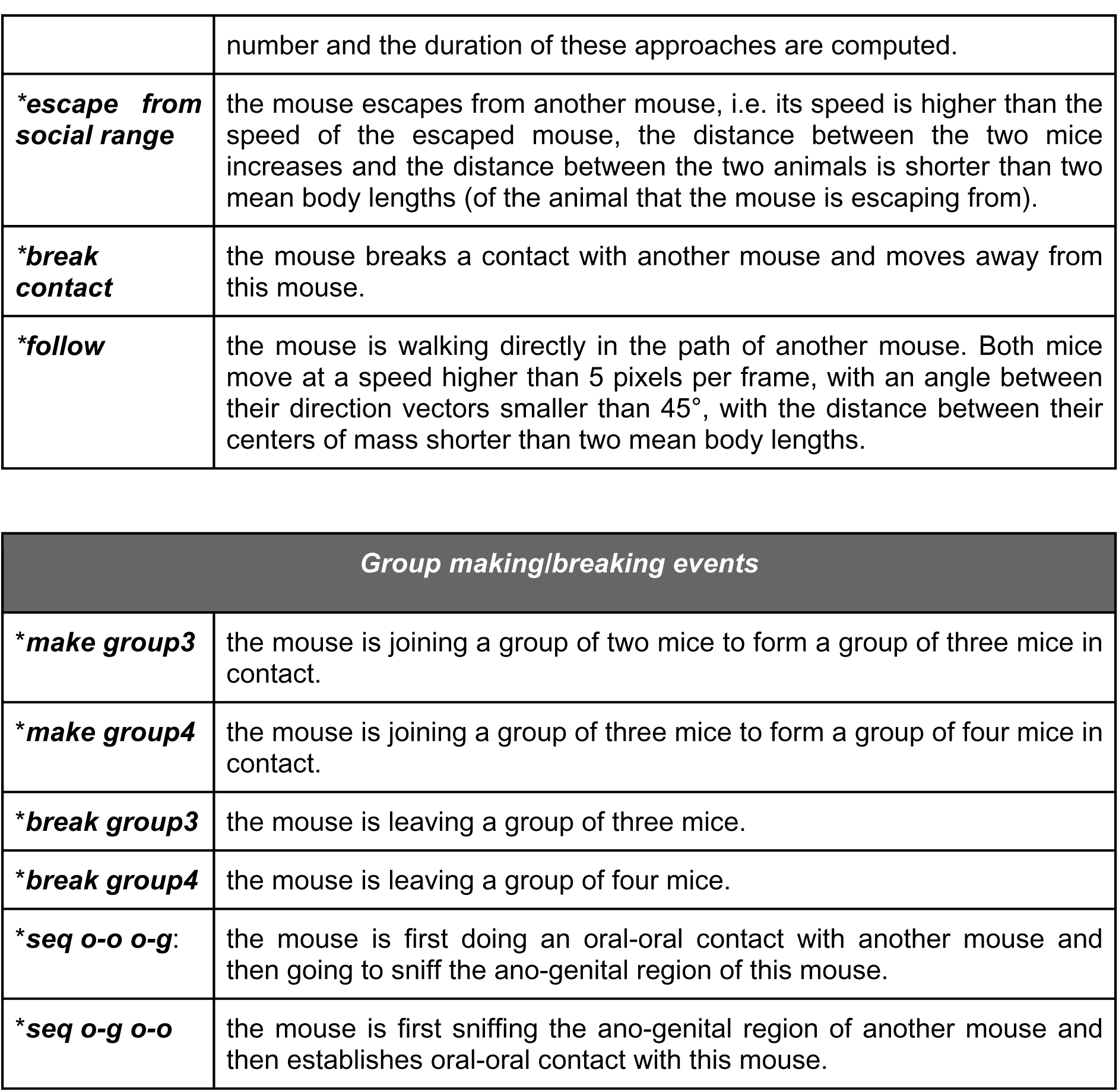

### Dynamics of subgroups of three mice within groups of four mice

We aimed at studying the dynamic of groups. We detail here the formation and breaking of groups of three mice. We defined the joiner and the breaker of these groups by identifying the mouse approaching and getting into contact or breaking contact at the frame just preceding or just following the group3 event, respectively. In the following paragraph, we detail how we calculate the odds of these events.

We consider only the initiation of a group of three mice (but the reasoning is similar for the ending of a group of three mice). With two wild-type (WT_1_, WT_2_) and two mutant (KO_1_, KO_2_) mice initially in the cage, six pairs are possible: WT_1_-WT_2_, KO_1_-KO_2_, WT_1_-KO_1_, WT_1_-KO_2_, WT_2_-KO_1_ and WT_2_-KO_2_.

When calculating the expected probability of who is joining two mice to form a group of three mice, the odds to have an initial pair including two wild-type mice are 1/6. The probability of a mutant mouse to join this pair in 1/1 since it is the only genotype available. Therefore, the chance to have a mutant mouse joining a pair of wild-type mice is p(KO|WT-WT) = 1/6 × 1/1 = 1/6. With a similar reasoning:

-The odds to have an initial pair including two mutant mice are 1/6. The probability of a wild-type mouse to join this pair in 1/1 since it is the only genotype available. Therefore, the chance to have a wild-type mouse joining a pair of mutant mice is p(WT|KO-KO) = 1/6 × 1/1 = 1/6.
-The odds to have an initial pair including one wild-type mouse and one mutant mouse are 4/6. The probability of a wild-type mouse to join this pair is 1/2 since a wild-type and a mutant mouse are available. Therefore, the chance to have a wild-type mouse joining a pair of one wild-type mouse and one mutant mouse is p(KO|WT-KO) = 4/6 × 1/2 = 1/3.
-The odds to have an initial pair including one wild-type mouse and one mutant mouse are 4/6. The probability of a mutant mouse to join this pair is 1/2 since a wild-type and a mutant mouse are left. Therefore, the chance to have a mutant mouse joining a pair of one wild-type mouse and one mutant mouse is p(WT|WT-KO) = 4/6 × 1/2 = 1/3.

These probabilities can also be found by counting the possible groups (initial pair - joiner or breaker):

|  |  |  |  |
| --- | --- | --- | --- |
| WT<sub>1</sub>,WT<sub>2</sub> - KO<sub>1</sub> | WT<sub>1</sub>,WT<sub>2</sub> - KO<sub>2</sub> |  |  |
| KO<sub>1</sub>,KO<sub>2</sub> - WT<sub>1</sub> | KO<sub>1</sub>,KO<sub>2</sub> - WT<sub>2</sub> |  |  |
| WT<sub>1</sub>,KO<sub>1</sub> - WT<sub>2</sub> | WT<sub>1</sub>,KO<sub>1</sub> - KO<sub>2</sub> | WT<sub>1</sub>,KO<sub>2</sub> - WT<sub>2</sub> | WT<sub>1</sub>,KO<sub>2</sub> - KO<sub>1</sub> |
| WT<sub>2</sub>,KO<sub>1</sub> - WT<sub>1</sub> | WT<sub>2</sub>,KO<sub>1</sub> - KO<sub>2</sub> | WT<sub>2</sub>,KO<sub>2</sub> - WT<sub>1</sub> | WT<sub>2</sub>,KO<sub>2</sub> - KO<sub>1</sub> |

### MP4 video recording

To be able to go back to visual data for control, we record the infrared stream in MPEG. We add overlay information to display tracking information, RFID state, time of acquisition, memory load, CPU time computation for the last frame and real time/date. MPEG is recorded in full infrared resolution (512 × 424) at half the input rate by default (15 fps), customizable by user. MPEG is recorded in its own thread in low priority, meaning that in weak configuration, frames will be dropped from record. For storage and browsing convenience, each MPEG is 10 minutes long.

Code: *package livemousetracker.MPEGRecorder.*

### Database player

The database player allows to read database by replaying the data extracted from the software. Its GUI displays color-coded animal segmentation and the event corresponding to the current time points. Events can be displayed, no matter if they were computed from the original tracker, or subsequently in R, matlab, python or any software. If events are related to only one animal, the segmentation carries those descriptions. Otherwise, a column showing all interaction is color coded to better understand which mice are involved. Events can be filtered to help studying specific events. Users can seek forward or backward in the database with different time-steps, or change the frame rate. code: *package livemousetracker.dataplayer.* **Supplementary movie: LMT - Method overview**.

### Collaborative website sharing data

The website is opened to anybody (**Supplementary Fig. S7**). Users first store their data in the location they prefer, and then they reference their datasets within this website. People can then search for experiment and download databases, films, script to process data and any material users want to share.

**Supplementary Fig. S7.**
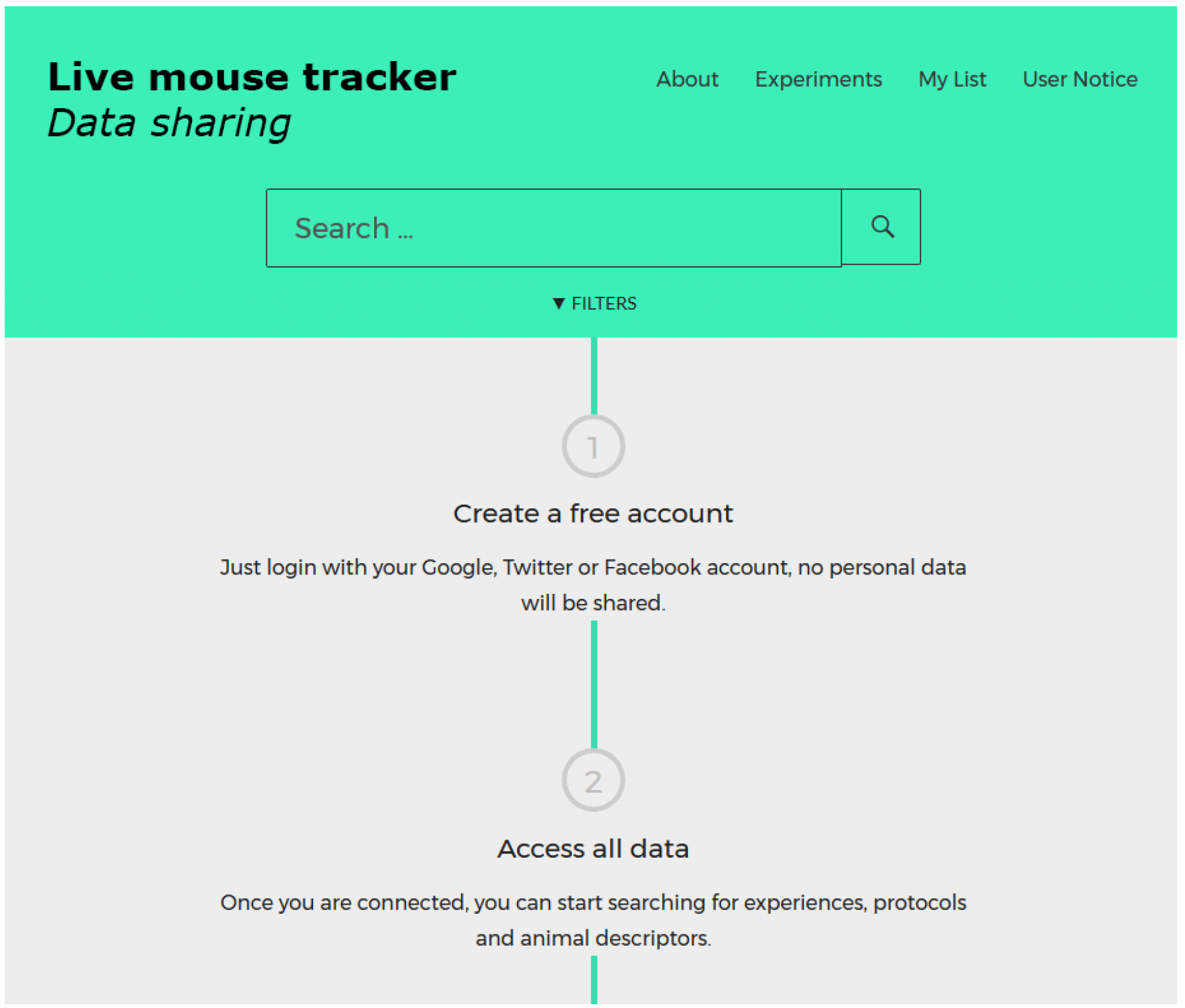
Opening view of the website http://livemousetracker.org designed to share experiments.

### Software Development Kit

This work represents two years of work dedicated to the development of this complete solution. A large part of the solution consists in connecting to driver, grabbing stream, ensuring that streams are synchronized, dealing with java real time consideration, creating code probe to help debugging, creating display overlay, creating streamed data structure, dealing with calibration, output of data as video stream, database or network stream for immediate control of third party devices. For developing purposes, a number of side tools have been created, that are only used by the developers, not by the users. They consist in small programs that aim at helping the developers to debug code and test new features. Those tools are also provided in open source to build on top of the existing tracking. Among those tools, developers will find a raw data recorder and player, which mimics a live kinect device, but allows to play several times the exact same film.

A large part of those developments is not to said “scientific”, but it is mandatory to build a final working product. We documented all this trunk of code so that newcomer can test their own techniques directly in this framework, in order to focus their energy on the problem of enhancing the phenotyping instead of working on all the side engineering problems.

Finally, to ease distribution of new version, Live Mouse Tracker is developed as a plugin of Icy, which makes easy fork and further distributions.

### Mice

*ProSAP1/Shank2* mutant mice (hereafter named *Shank2* mutant mice) were initially described in a previous study^8^. They were bred on a C57Bl/6J background (>12 backcrosses), and maintained by crossing heterozygous parents. We tested adult female littermates aged between 3 and 12 months of age (the experiments were spread over several months). For the single object exploration, we used 10 *Shank2*^+/+^ and 8 *Shank2*^−/−^ female mice aged of 4 months in the first cohort. These mice also underwent a previous behavioral characterization with classical methods (data not shown). The second cohort included 12 *Shank2*^+/+^ and 7 *Shank2*^−/−^ female mice and 8 *Shank2*^+/+^ and 8 *Shank2*^−/−^ male mice (aged of 2.5-4 months at the time of testing). For the object exploration in pairs, we used 12 *Shank2*^+/+^ and 12 *Shank2*^−/−^ mice aged of 9 months at the time of testing in the first cohort (only two *Shank2*^+/+^ and four *Shank2*^−/−^ did not undergo the single object exploration test). We used 13 *Shank2*^+/+^ and 11 *Shank2*^−/−^ female mice aged of 5-6 months from the second cohort (all of them underwent the single object exploration task at least three weeks before).

*ProSAP2/Shank3* mutant mice (hereafter named *Shank3* mutant mice) were initially described in a previous study^8^. They were bred on a C57Bl/6J background (>10 backcrosses) and maintained by crossing heterozygous parents. We tested adult female littermates (from heterozygous parents). For the single object exploration, we used 7 *Shank3*^+/+^ and 12 *Shank3*^−/−^ female mice aged of 2.5-3 months. We did not conduct paired object exploration with the *Shank3* mouse strain.

For the group behavior experiment in four animals per cage, we constituted social groups at least three weeks before the experiments. We focused our study on female mice. Indeed, constituting mixed-genotype groups of four mice after weaning was not possible with males given their aggressiveness toward unfamiliar same-sex conspecifics at maturity^8^. We constituted nine groups of four mice over the two cohorts of *Shank2* mutant mice (two *Shank2*^+/+^ and two *Shank2*^−/−^), which represented 18 *Shank2*^+/+^ and 18 *Shank2*^−/−^ mice aged of 3-13 months at the time of testing (all of them except two *Shank2*^+/+^ and two *Shank2*^−/−^ performed the previous experiments). We also constituted six groups of four mice from the *Shank3* strain, with two *Shank3*^+/+^ and two *ShankS*^−/−^ mice, which represented 12 *Shank3*^+/+^ and 12 *Shank3*^−/−^ female mice. These mice were aged of 3 to 4 months at the time of testing. To be able to compare the two models, we also tested only adult females in the *Shank3* strain. For both strains, the ages were homogenous within each group of four mice to get age-matched mutant and control mice

At least 3 weeks separated two consecutive experiments. Mice were housed in standard laboratory cages in same-sex littermate groups of two to four mice until the single object exploration. After the single object exploration, for paired object exploration, mice were housed in pairs (wild-type/wild-type, wild-type/mutant and mutant/mutant; mixing non littermate mice) at least one week before the experiments. Finally, groups of four mice were constituted at least three weeks before the group monitoring study and were not changed anymore. Overall, social groups were changed maximum three times over the course of the experiment pipeline. *Shank2* mice were housed in a 12:12 dark-light cycle, with lights on at 07:00 AM, while *Shank3* mice were housed in a 11:13 dark-light cycle, with lights on at 07:00 AM. Food and water were available ad libitum.

Mice of both strains were identified at weaning (four weeks of age) using ear punches. The skin sample was used for genotyping following the protocols described in the original publication^8^. Between two and three months of age, we inserted the RFID tag subcutaneously under isoflurane anesthetic with local analgesia (lidocaine<0.05 ml at 21.33 mg/ml). All experiments involving animals complied with the European ethical regulation, and were validated by the ethical committee CETEA n°89, Institut Pasteur, Paris.

### Behavioral protocols

#### Single and paired object exploration

We placed the experimental cage (50×50×30 cm) under the setup (70 lux, T=22°C). New fresh bedding (2-3 cm high) covered the bottom of the cage; bedding is renewed for each animal. We placed the tested mouse in the test cage and left it to freely explore the apparatus for 30 min (phase 1). After these 30 min of free exploration, we introduced a novel object (red Plexiglas house, permeable to infra-red light; 9.5×7.5×4.5 cm; Special Diet Services, England) in the bottom left quarter of the cage. The mouse was left for 30 min in the apparatus (phase 2). In the paired condition of the first cohort, we constituted three types of pairs (non littermates animals): 4 pairs of *Shank2*^+/+^ & *Shank2*^+/+^, 4 pairs of *Shank2*^+/+^ & *Shank2*^−/−^ and 4 pairs of *Shank2*^−/−^ & Shank2^−/−^. In the second cohort, we constituted 4 pairs of *Shank2*^+/+^ & *Shank2*^+/+^, 4 pairs of *Shank2*^+/+^ & *Shank2*^−/−^ and 4 pairs of *Shank2*^−/−^ & *Shank2*^−/−^. These pairs were together for at least one week before the experiment. We used the software Python 3.6 (Python Software Foundation. Python Language Reference, version 3.6; available at http://www.python.org) to compute distances and stretched attend posture in the experiments.

#### Long-term monitoring of groups of four mice

Nine groups of four mice were constituted with *Shank2* mice and six groups of four mice were constituted with *Shank3* mice. We grouped together two wild-type and two mutant mice at least three weeks before the experiment. We do not control the fact that wild-type mice may be influenced by the behavior of the two mutant mice; however we consider these variations as within the normal variations within a population and analyzed therefore the data at the individual level. Each group of four mice was placed in the test cage (50×50×30 cm; 70 lux when the light was on and 0 lux when the light was off; T=22°C), with one red house (see above in the single object exploration test), 6 cylindrical compressed cottons, and food and water ad libitum. Recording started immediately for 23 h for the *Shank2* mice and for three days for the *Shank3* mice. We used the software Python 3.6 (Python Software Foundation. Python Language Reference, version 3.6; available at http://www.python.org) to compute distances and behavioral events from the database.

#### Statistical analyses

We used the software R 3.3.1 for graphical representation and statistics (R Core Team (2016). R: A language and environment for statistical computing. R Foundation for Statistical Computing, Vienna, Austria. URL https://www.R-project.org/). For the single and dyadic social interactions, we used non-parametric Wilcoxon rank sum tests to compare variables between genotypes, given the non-normality of the data and the small sample size. For the long-term recordings of groups of four mice, we used non-parametric Wilcoxon rank sum tests to compare behavioral events occurrences and duration between genotypes, as well as t-tests to compare the dynamic of the group formation. When necessary, we applied Bonferroni corrections for multiple testing.

### Software and system general description

We developed the methods as a plugin of Icy. The code is developed in Java 8 (189 classes. 19934 lines of code. 950 Methods. 32 Packages.). We put specific effort to make the code readable and documented for further extension. Experiments were first conducted on intel core i5-6400/2.4GHz/16GB and then on Ryzen 1800×. We recommend this last version.

### Motivation and review of the existing tracking methods

We present here an overview of the current tracking methods used to follow mice and to extract behavioral information. We selected here the most representative ones of each type of method used. We first present the current needs in phenotyping studies and then we detail to what extent the different methods answer these needs.

Phenotyping mouse models usually involves one individual screened through various behavioral tests to measure anxiety, learning, memory, or locomotion. Social aspects are determined in dyadic interactions, but these aspects are the least examined and remain to be complemented in existing phenotyping databases^27,28^. With the aim of documenting in the most comprehensive way the phenotype of an individual, studying social aspects within a group of animals would allow to document both dyadic interactions and more complex ones involving more than one other individual. These two aspects of sociality can therefore be dissociated and will enrich phenotyping data^19,29–32^. Such approaches will increase the translational values of the phenotyping studies^24,33^.

In clinical examination of patients with neuropsychiatric disorders affecting social relationships, both the initiation and the maintenance of these relationships are evaluated (ADI-ADOS for autism spectrum disorders). Therefore in mouse models for these disorders, we should also take into account these aspects by following the freely moving animals over longer durations^24^. Indeed, short experiments have been shown to be misleading in some cases^34^. New experiments should be built to gather precise data on individual mice over the long term to answer the more and more precise needs of phenotyping with ethologically-relevant measures^27,28^.

Individual following is necessary to identify individual profiles. This is required to model personalized medicine. However, visual or olfactory marks could lead to modifications of the general behavior of the animals (as in zebrafishes^35^), biasing the results of the experiment. Therefore, individual following should be conducted without perceptible marks on the animals.

To summarize these requirements, the current need in phenotyping is to measure automatically in a detailed way social and non-social behaviors of individuals within a group. The description of complex behaviors requires a high level of details. An attempt was already made to reach a high level of details in body postures with a depth sensor camera but this was used on an isolated mouse in an openfield^7^. A high level of details in social contexts is currently reached when only two individuals are tested together over short durations (i.e., 10-15 min). The way such data is collected does not allow users to investigate more individuals simultaneously or over a longer duration. de Chaumont and colleagues (2012) published a semi-automatic software to describe precisely the types of interactions and the sequences of social events occurring between two visually indistinguishable mice, from a top-view video camera^36^, but this solution is limited as it need to be supervised and corrected by an expert. Unger and colleagues (2017) built on MiceProfiler and used another segmentation method to improve detection and avoid manual corrections^37^ (manual interventions remained necessary at the beginning of the tracking). Hong and colleagues (2013) developed a system with two video cameras and one depth sensor to follow two mice^38^, but they needed to be of different coat colors to track the identities. The data extracted were detailed (attack, mounting, distances head to body and body to body), but they were calculated based on annotated video frames and training of a classifier. Finally, the highest level of details was provided by Matsumoto and colleagues (2013). They developed a system using four depth sensors to follow in details the different steps of mating in rats in full 3D over ten minutes^39^. It only requires manual intervention to define the initial position of the models. Giancardo and colleagues (2013) have developed a system able to track unmarked mice over one hour^40^. The behavioral repertoire measured was large (types of contacts, above, following, non-social behaviors such as walking or standing alone), but the system needed an average of 1 manual intervention to correct identities every 34 s (in a 3-mice experiment). Overall, all these systems need manual interventions to provide the most relevant data. This is manageable over short experiments up to one hour, but not over longer experiments of several hours or days.

For such long-term experiments, the possibilities to track individually animals in groups are two folds. First, the animals are marked visually, using hair dye. These visual marks can be combined with radio frequency identification (RFID) tags or not. So et al. (2015) manually scan sampled over 21 days chasing, fighting, sniffing and allo-grooming^29^. This approach provided impressive data on hierarchy and social relationships within the group, but it is not manageable on long-term experiments in mass phenotyping. Similarly, in the Visible Burrow System^41^, Bove and colleagues (2018) manually annotated six 10-min periods per day over five days of a group of eight mice, either from the C57BL/6J strain or from the BTBR strain and highlighted a reduced level of social activity and an increased level of grooming, but no difference in environmental exploration, social avoidance or aggression (except on the first day)^33^. Howerton et al. (2012) scored the activity of two mice and their grouping within compartments automatically over 24 hours^42^. They used the reads of RFID tags when mice travelled through antennas and combined data with visual validation using hair dye. Similarly, Shemesh et al. (2013) proposed a system providing the activity as well as the spatial occupancy of the animals marked visually, whose identity was learned over labeled video frames^30^. Castelhano-Carlos et al. (2014) conducted the same kind of activity and position data on rats identified by RFID tags and pen marks on the tail^43^. Interestingly, they also enriched their data with social behaviors that were annotated manually. The most advanced system in this category with animals visually marked is the system of Ohayon and colleagues (2013). They provided data over five days on the place preference of the mice, as well as their social grouping and following behaviors (based on pre-labeled videos)^19^. Altogether, these methods using visual marks to track animals (verified through RFID tag scanning or not) provided most of the time activity and place preference data. They are based on contrast and need specific fixed experimental conditions, obtained through background simplification or infrared lightning. Additional social behaviors needed to be manually annotated, except in the system of Ohayon et al. (2013)^19^.

For this second set of methods, the animals are not visually marked. In most of these systems, the animals are identified using RFID tags, and the only data extracted are on the activity and the spatial grouping over different compartments/zones of the cage^32,44–46^. This method provides the longest experiment possible, but with no precise social data. Perez-Escudero and colleagues (2014) developed a very interesting and inspiring method to identify each individual based on a visual fingerprint^47^. This digital signature is coded by associating contrast information and distances within the animal. This system, idTracker, works very nicely with fishes and ants, as these animals are geometrically incompressible: indeed, even if they visually bend, they still always display the same surface. Mice in contrast are highly deformable, able to stretch or contract with a large spectrum of different conformations, eventually displaying different fur appearance, and, in addition, they can stand up (known as rearing) and present yet another set of visual conformations. This makes IdTracker unable to keep the identity of mice because the geometrical/digital signature used cannot deal with such a deformable model. The level of details provided by this method is not sufficient to characterize globally the social behavior of mouse models of neuropsychiatric disorders. With such a system, Perez-Escudero and colleagues managed to extract spatial occupancy, velocity or relative position of the animals within a group of zebrafishes, but no robust data on mice. Weissbrod and colleagues (2013) combined RFID identification with video recordings and provided social data reflecting relative movements, with a reduced behavioral repertoire but no robust data on the contacts between animals^31^. Finally, Alexandrov and colleagues (2015) proposed a system that can supposedly provide activity and contact data between non-visually marked animals but no information on the method used and on the validity of the tracking is available^48^. RFID tags are the most robust way to identify individuals but they have their own constraints in reading speed and in probe range. The RFID tag should be 1/ in the probe range for 1/10th s, 2/ the only tag in this range, and 3/ read by only one antenna, meaning only one antenna should be activated at a time. These constraints on RFID explain why their usage is not trivial, for instance, RFID floor, sold by the TSE company (TSE Systems GmbH European / Asian headquarters, Bad Homburg, Germany), cycle blindly over the antenna to read RFID, therefore the probability of reading an animal is reduced again by the number of antenna integrated in this device.

All these methods have their own advantages, but none of them provide the possibility of track mice individually within a group over long-term experiments with the level of details that is requested for phenotyping mouse models of neuropsychiatric disorders. In addition, none of these systems is real time and therefore there is no possibility to directly interact with the experiment, despite the fact that several groups already asked for this feature^38,39^. This would allow to increase phenotyping speed^42^, and reject all possibility of manually correcting of the data. Tracking errors should therefore be managed directly at the source, and not corrected afterwards. As fulfilling these needs with the current methods of animal tracking is not possible, we developed our own system working in real time and managing identification errors.

### Supplementary results

**Supplementary Fig. S8.**
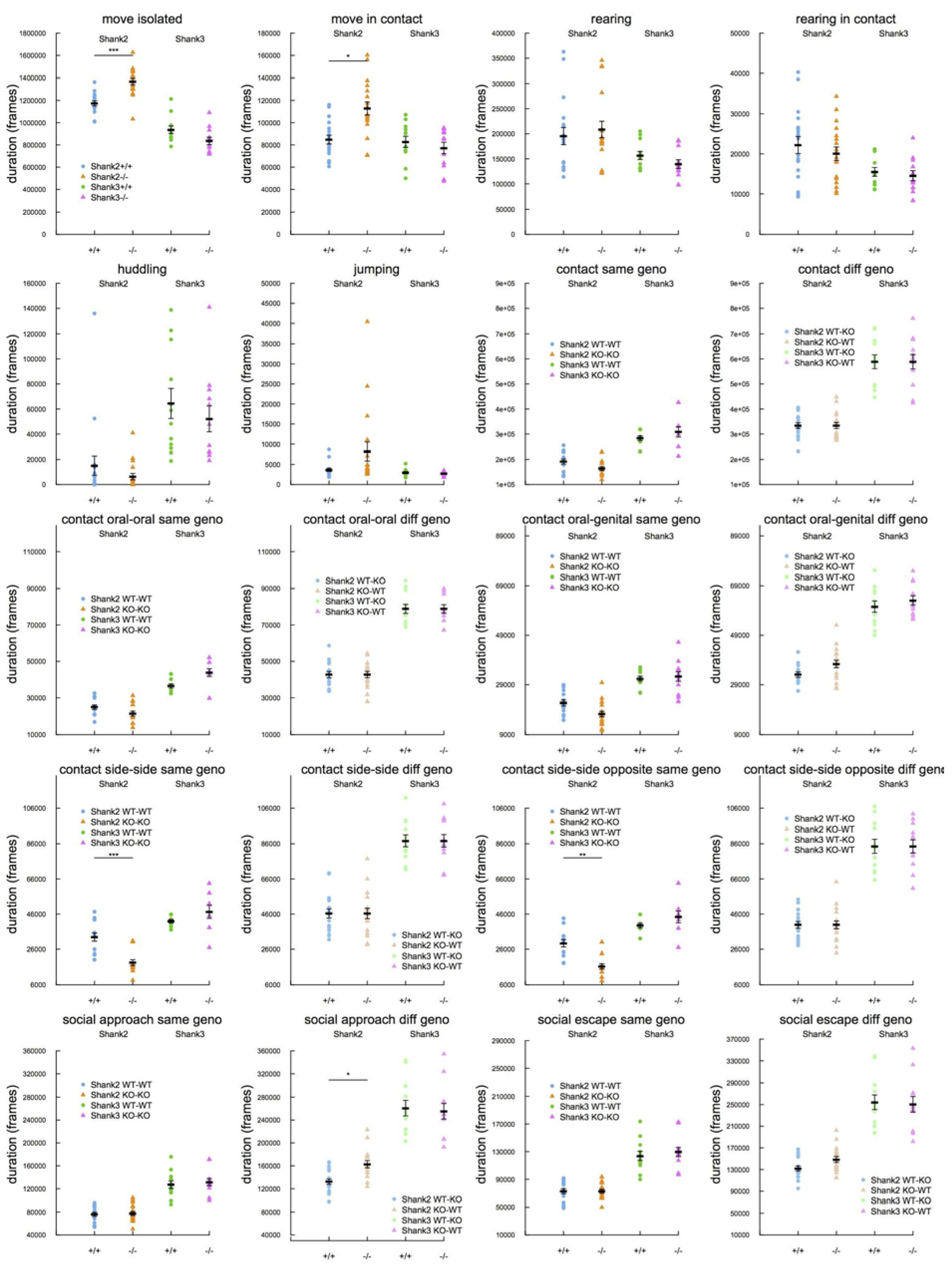
Total duration of a subsample of the behavioral events extracted from the monitoring over 23h of *Shank2* and *Shank3* mixed-genotype groups of four mice (Wilcoxon rank sum test; *: p<0.05; **: p<0.01; ***: p<0.001).

**Supplementary Fig. S9.**
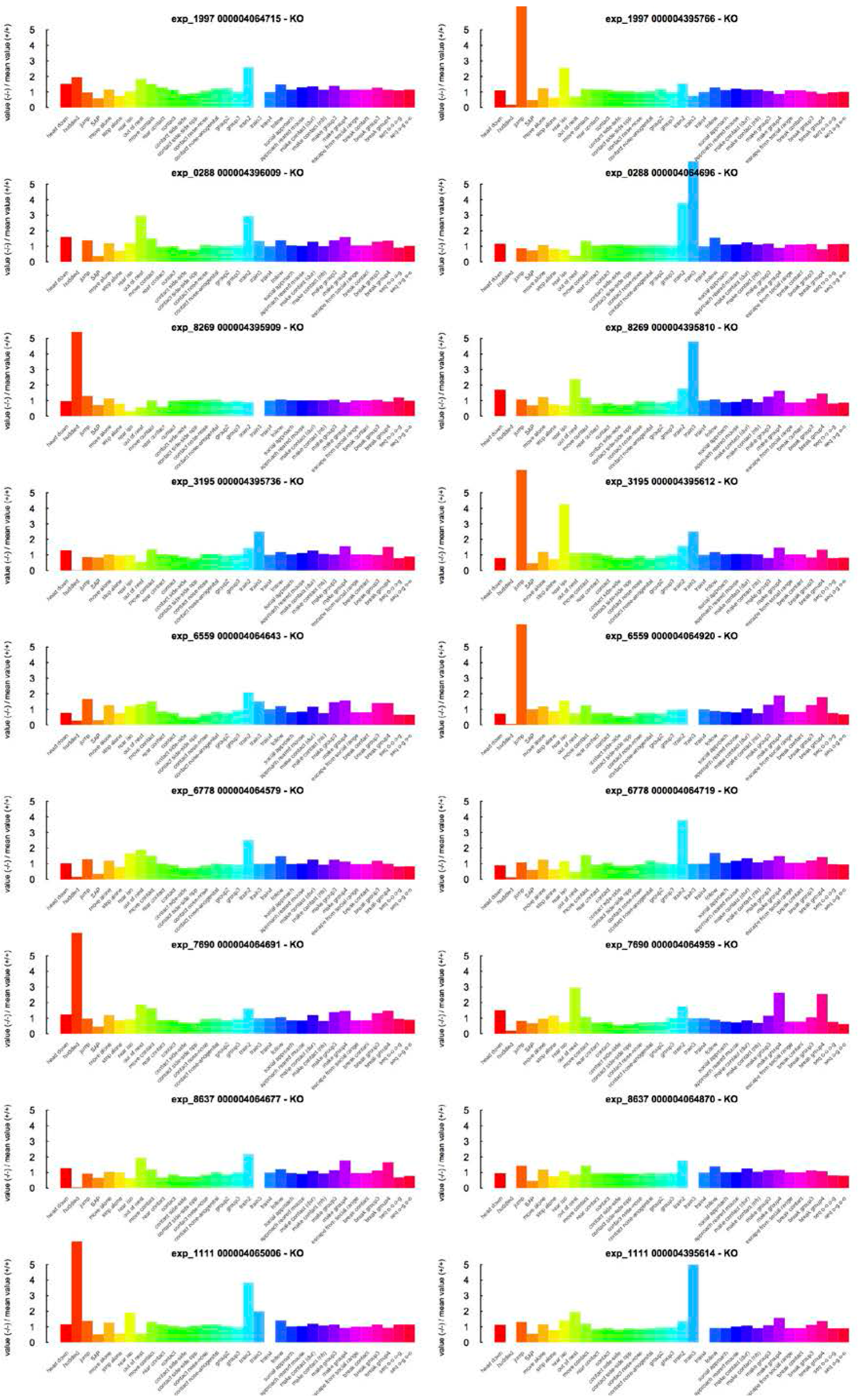
Individual profiles of each *Shank2^−/−^* mouse calculated over 23h of recording. One row represents the profiles of each of the two mutant mice from the same experiment. Traits that were not different from the mean value of the wild-type mice of the experiment were set at zero. Traits that are more expressed in mutant mice than in the mean of wild-type mice have positive values, while traits that are less expressed in mutant mice than in the mean of wild-type mice have negative values. Y-axis graduations represent the order of magnitude of the difference between mutant and wild-type mice, i.e., a value of 2 represents a trait that is two times more expressed in the mutant mice than in the wild-type mean mouse.

**Supplementary Fig. S10.**
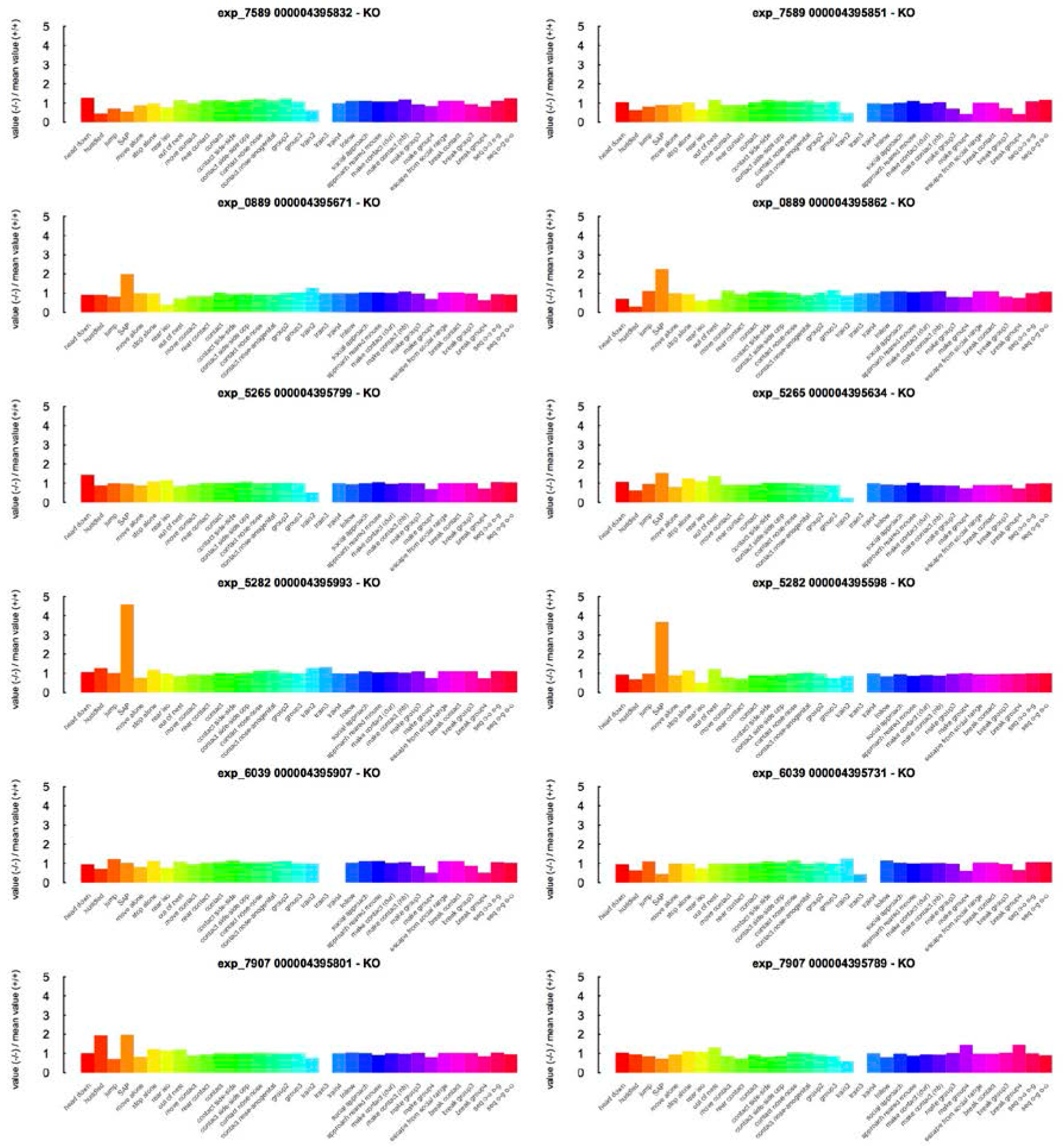
Individual profiles of each *Shank3*^−/−^ mouse calculated over 23h
recording. One row represents the profiles of each of the two mutant mice from the same experiment. Traits that were not different from the mean value of the wild-type mice of the experiment were set at zero. Traits that are more expressed in mutant mice than in the mean of wild-type mice have positive values, while traits that are less expressed in mutant mice than in the mean of wild-type mice have negative values. Y-axis graduations represent the order of magnitude of the difference between mutant and wild-type mice, i.e., a value of 2 represents a trait that is two times more expressed in the mutant mice than in the wild-type mean mouse.

**Supplementary Fig. S11.**
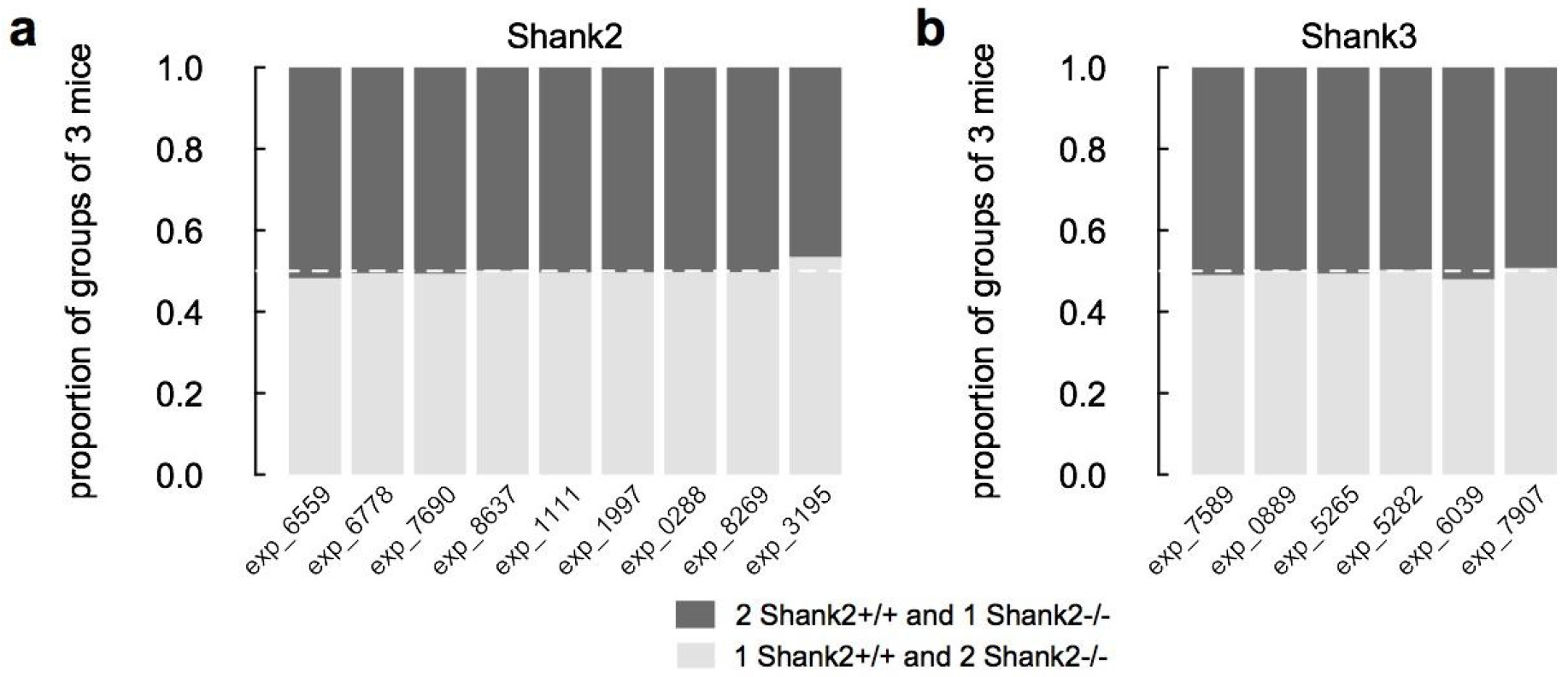
Proportions of groups of three individuals for each one-day recording of four individuals for (a) *Shank2* mice and (b) *Shank3* mice. Two types of groups of three individuals were possible within the groups of four mice: two mutant mice with one wild-type mouse (light grey) and one mutant mouse with two wild-type mice (dark grey). The expected proportion (white line) was 0.5 given the composition of the cage (two mutant mice and two wild-type mice). We used a Chi square conformity test. No significant differences were found.

**Supplementary Fig. S12.**
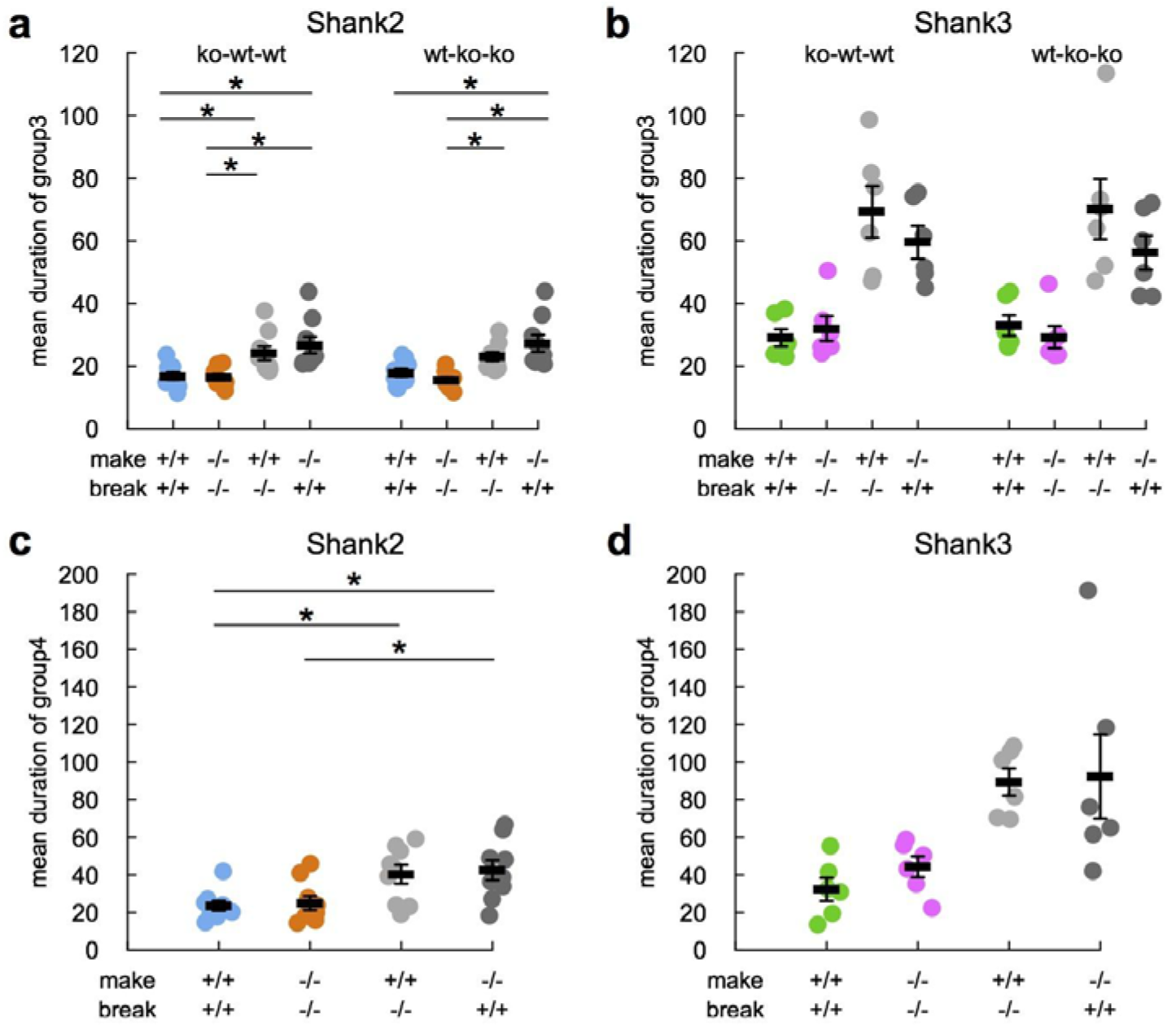
Mean duration (frames) of groups of three and four individuals for each one-day recording of four individuals for (a) *Shank2* mice and (b) *Shank3* mice according to who is coming in and coming out. Wilcoxon tests with Bonferroni corrections for multiple testing. Data are present as mean±SEM. *Shank2* strain: nine experimental groups; *Shank3* strain: six experimental groups.

**Supplementary Fig. S13.**
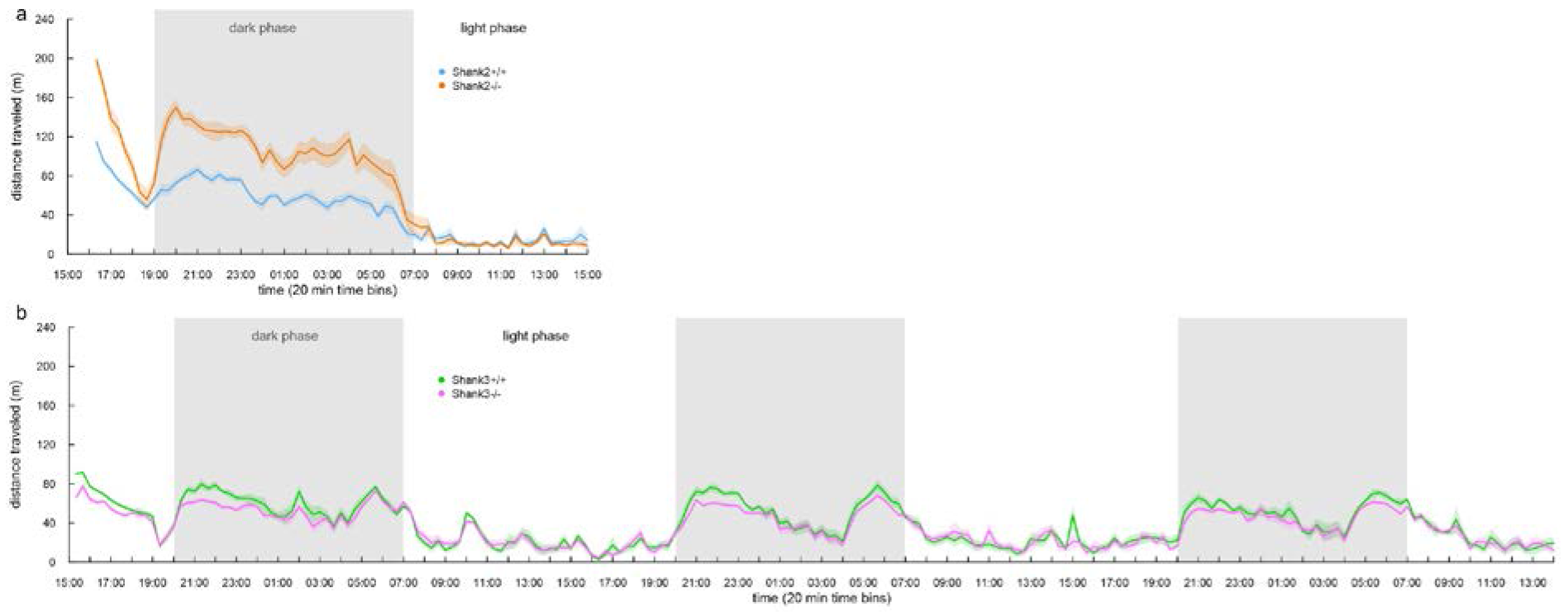
Mean activity levels of (a) *Shank2* mice and (b) *Shank3* mice over one day and three days of recording, respectively. We computed the distance traveled by each mouse per 20 min time bins. We present the mean distance traveled per time bin for each genotype. *Shank2* strain: 18 *Shank2*^−/−^ mice & 18 *Shank2*^+/+^ mice; *Shank3* strain: 12 *Shank3*^−/−^ mice & 12 *Shank3*^+/+^ mice.

**Supplementary Fig. S14.**
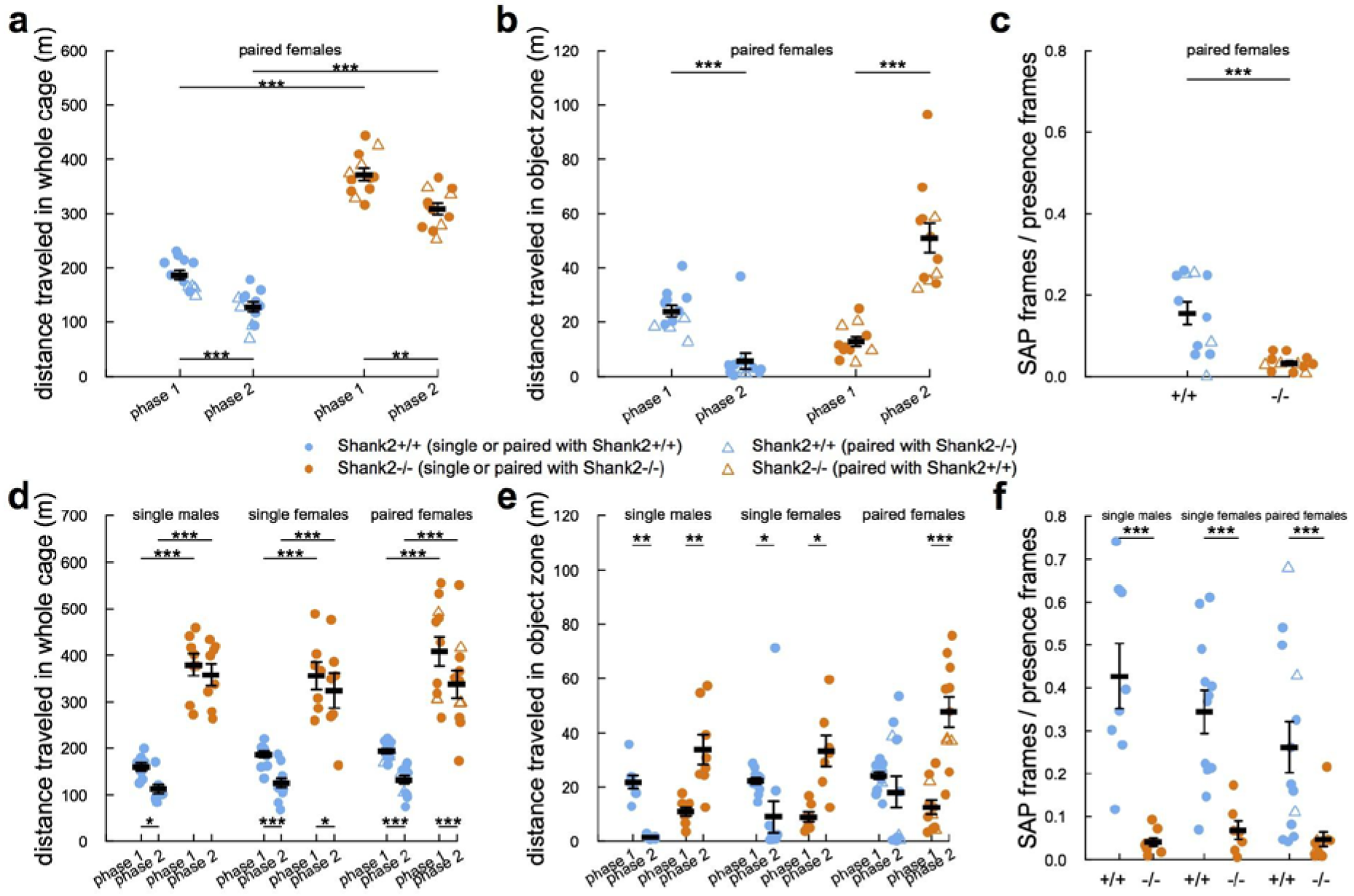
Hyperactivity and atypical exploration strategy in paired conditions and confirmed in a second cohort of *Shank2*^−/−^ mice. **(a)** Distance traveled in the entire cage in phase 1 and phase 2 for wild-type and *Shank2*^−/−^ female mice from cohort 1 in paired conditions. **(b)** Distance traveled in the object zone in phase 1 and phase 2 for wild-type and *Shank2*^-1-^ female mice from cohort 1 in paired conditions. **(c)** Ratio of the number of frames in the object zone where the animal was detected in stretched attend posture over the total number of frames spent in the object zone. **(d)** Distance traveled in the entire cage in phase 1 and phase 2 for wild-type and *Shank2*^−/−^ mice from cohort 2 for males in the single condition, and for females in the single and paired conditions. **(e)** Distance traveled in the object zone in wild-type mice and *Shank2*^−/−^ mice from cohort 2 in phase 1 and phase 2 for males in the single condition, and for females in the single and paired conditions. **(f)** Proportion of frames in which mice are in SAP in the object zone over the total number of frames in which animals are detected in the object zone. Data are presented as mean±sem and individual points for: (a-c) cohort 1: 12 *Shank2*^+/+^ mice and 12 *Shank2*^−/−^ mice; (d-e) cohort 2: 12-13 *Shank2*^+/+^ female mice and 7-11 *Shank2*^−/−^ female mice and 8 *Shank2*^+/+^ male mice and 8 *Shank2*^−/−^ male mice in cohort 2 (Wilcoxon rank sum test; *: p<0.05; **: p<0.01; ***: p<0.001).

